# Pharmacological inhibition of longevity regulator PAPP-A restrains mesenchymal stromal cell activity

**DOI:** 10.1101/2020.02.05.936310

**Authors:** Mary Mohrin, Justin Liu, Jose Zavala-Solorio, Sakshi Bhargava, John Maxwell Trumble, Alyssa Brito, Dorothy Hu, Daniel Brooks, Mary L. Bouxsein, Roland Baron, Yuliya Kutskova, Adam Freund

## Abstract

Reducing insulin-like growth factor (IGF) signaling is one of the best conserved and characterized mechanisms to extend longevity. Pregnancy associated plasma protein A (PAPP-A) is a secreted metalloprotease that increases IGF availability by cleaving IGF binding proteins. PAPP-A inhibition reduces local IGF signaling, limits the progression of multiple age-related diseases, and extends lifespan, but the mechanisms behind these pleiotropic effects remains unknown. Here, we developed and utilized a PAPP-A neutralizing antibody to discover that adulthood inhibition of this protease reduced collagen and extracellular matrix (ECM) gene expression in multiple tissues in mice. Using bone marrow to explore this effect, we identified mesenchymal stromal cells (MSCs) as the source of PAPP-A and primary responders to PAPP-A inhibition. Short-term treatment with anti-PAPP-A reduced IGF signaling in MSCs, altered MSC expression of collagen/ECM, and decreased MSC number. This affected MSC-dependent functions, decreasing myelopoiesis and osteogenesis. Our data demonstrate that PAPP-A inhibition reduces the activity and number of IGF-dependent mesenchymal progenitor cells and their differentiated progeny, and that this reduction leads to functional changes at the tissue level. MSC-like cells are present in virtually all tissues, and aberrant collagen and ECM production from mesenchymal cells drives aspects of aging and age-related diseases, thus this may be a mechanism by which PAPP-A deficiency enhances lifespan and healthspan.

**Summary:** Inhibition of PAPP-A, a regulator of IGF signaling, decreases multi-tissue collagen and extracellular matrix gene expression and modulates mesenchymal stromal cell activity in murine bone marrow.

## Introduction

Initially appreciated for its essential role in fetal and postnatal development and growth, the insulin-like growth factor (IGF) signaling pathway has also been implicated in the regulation of aging and lifespan since the discovery that partial loss-of-function mutations in the C. *elegans daf-2* gene, which encodes an ortholog of the human insulin and IGF receptors, can double lifespan *(1)*. Since this discovery, a reduction in IGF signaling has been found to delay aging and extend lifespan in multiple species, including mammals *(2–4)*. A number of mutations in the IGF signaling pathway have been reported to extend lifespan and delay aging in the laboratory mouse, but therapies to target IGF signaling for age-related diseases, let alone aging itself, remain absent from the clinic *(5)*. Clinical trial failures of IGF receptor inhibitors for various malignancies have led some to conclude that targeting the core components of this pleiotropic pathway (e.g., IGF-1 and IGF-1R) is a therapeutic dead-end, largely due to the existence of complex compensatory signaling from growth hormone, insulin, and other growth factors *(6–8)*.

Pregnancy associated plasma protein A (PAPP-A) is a metalloprotease with the ability to cleave IGF binding proteins (IGFBPs), specifically IGFBP-2, −4, and −5, and provides a way to tune IGF signaling that circumvents some of the problems encountered by IGF-1R inhibitors *(9)*. Altering PAPP-A activity does not affect the systemic IGF-1 circuit: circulating levels of GH and IGF-1 do not change, and thus do not elicit compensatory feedback or elevation of GH signaling via the somatotropic axis *(10, 11)*. By cleaving IGFBP-2, −4, or −5 and liberating IGFs to signal through their receptor, PAPP-A increases local IGF signaling; conversely, inhibition of PAPP-A allows IGFBPs to accumulate, blocking IGFs from binding their receptors and reducing IGF signaling. Although PAPP-A is secreted, it is localized to the cell surface by binding to glycosaminoglycans *(12)*, perhaps explaining why its effects are primarily local.

The physiologically relevant role of PAPP-A in tuning IGF signaling is clear from the phenotypes of mice deficient for PAPP-A (PAPP-A KO). PAPP-A KO mice are proportional dwarfs, pointing to the role of IGF in growth modulation, and are also long-lived, with a ~30% increase in both male and female longevity: yet another piece of evidence supporting the role of IGF signaling in lifespan and aging *(10, 13)*. Importantly, constitutive deletion of PAPP-A in adulthood (at 5 months of age) also results in a substantial (21%) lifespan extension *(14)*, suggesting the mechanisms by which PAPP-A modulates longevity can be explored in the adult animal without confounding developmental effects - a characteristic that we have taken advantage of in this study. Importantly, the effects of PAPP-A inhibition on aging go beyond just lifespan extension; PAPP-A inhibition delays progression of age-related pathology in multiple tissues *(13)*, reduces fat accumulation on a high fat diet *(15, 16)*, delays age-related thymic atrophy *(17)*, delays atherosclerotic plaque progression and neointima formation *(18–20)*, and delays both spontaneous tumorigenesis and tumor progression in xenograft models *(13, 21–23)*. However, the mechanisms giving rise to this diverse set of phenotypes, or even the cell types affected by PAPP-A inhibition, remain largely unknown. Additionally, despite its importance in the regulation of growth and lifespan, the location and regulation of expression of PAPP-A remains relatively opaque. *Pappa* mRNA has been shown to have ubiquitous yet very low levels of expression across tissues with only modest changes during aging *(24)*. PAPP-A expression has been shown to increase after injury and during tissue remodeling; this regulation and its potential role in wound healing is not entirely understood *(25)*, though it may be related to inflammation: PAPP-A expression can be induced in response to inflammatory cytokines such as TNFa and IL-1β, and NF-κB binding motifs are present in the promoter of *PAPPA (11, 26, 27)*.

In short, attenuation of the GH/IGF axis extends longevity in multiple species and leads to the longest-lived mouse mutants known. Among these long-lived GH/IGF mutants, PAPP-A KO mice are unique in that they exhibit no feedback on the hypothalamic-pituitary axis. This highlights PAPP-A as an attractive target for therapeutic modulation of IGF signaling, but key aspects of PAPP-A biology are unknown: where is PAPP-A expressed? What tissues and cells are affected by the loss of PAPP-A? How does loss of PAPP-A extend health- and life-span? To address these questions, we generated a potent neutralizing antibody against murine PAPP-A and used it to discover that relatively short-term adulthood PAPP-A inhibition reduces collagen/ECM gene expression in multiple tissues. Exploring the mechanism behind this effect, we found that *Pappa* is highly expressed by mesenchymal stromal cells (MSCs), that PAPP-A inhibition reduces IGF signaling in MSCs and reduces their number, and that this results in multiple tissue-level consequences, affecting both myelopoiesis and osteogenesis. MSCs are important for normal tissue homeostasis, but with age and in the context of age-related disease, MSCs aberrantly expand and deposit excessive collagen and ECM *(28)*. Excessive collagen and ECM deposition are associated with aging in multiple tissues, and the reduced collagen/ECM production from MSCs and/or MSC-derived cells may be the mechanism by which PAPP-A inhibition delays aging and age-related disease in multiple tissues.

## Results

### Development of a potent and selective PAPP-A neutralizing antibody suitable for in vivo studies

PAPP-A has been reported to cleave IGFBP-2, IGFBP-4, and IGFBP-5 *(29–31)*. We tested this enzymatic activity biochemically, using recombinant components. As reported previously, PAPP-A demonstrated the ability to cleave all three binding proteins, and the enzymatic activity of PAPP-A for IGFBP-2 and IGFBP-4 was strongly enhanced by the addition of IGF-1 (Figure S1A). In contrast, the enzymatic activity of PAPP-A for IGFBP-5 was not increased by the addition of IGF-1 and was even slightly decreased, so for all subsequent experiments, cleavage of IGFBP-5 was tested in the absence of IGF-1. Also as reported previously, the activity of PAPP-A for each of these substrates varied, being the most active against IGFBP-5 and less active against IGFBP-2 and IGFBP-4 (Figure S1B).

As a specific IGFBP protease, PAPP-A is not thought to have IGF-independent effects, though there are isolated reports suggesting possible IGF-independent mechanisms *(32)*. To explore this, we tested whether PAPP-A alone induced gene expression changes in Hela cells. Hela cells express minimal endogenous *PAPPA* or *IGFBP4* compared to a cell type that expresses biologically-relevant amounts of *PAPPA*, such as IMR-90 fibroblasts *(30)* (Figure S1C), and we serum starved the cells overnight to remove any IGF-1 present in serum. Therefore, this was a useful system in which to study the effects of exogenous IGF-1, IGFBP-4, and PAPP-A on cell biology without endogenous interference. Addition of IGF-1 to the media of Hela cells stimulated a robust gene expression response, which was almost exactly recapitulated by the addition of IGF-1 + IGFBP-4 + PAPP-A (Figure S1D). In contrast, neither addition of PAPP-A nor addition of IGFBP-4 alone induced any change in gene expression. Although we cannot rule out the existence of IGF-independent effects of PAPP-A in other contexts or cell types, this result is consistent with neither PAPP-A nor IGFBP-4 having any biological function beyond modulating IGF signaling.

To study the functional role of PAPP-A in a variety of in vitro and in vivo contexts, we developed a neutralizing antibody against PAPP-A, which we call anti-PAPP-A. Anti-PAPP-A effectively blocked cleavage of both human and murine IGFBP-4 and IGFBP-2 by PAPP-A, but did not greatly affect PAPP-A’s ability to cleave IGFBP-5 (Figure S1E), similar to previously-generated PAPP-A neutralizing antibodies, and likely due to different requirements for cleavage between the IGFBP substrates *(29, 33–36)*, which remain to be better elucidated, e.g., by structural analysis. Anti-PAPP-A had sub-nanomolar affinity to murine PAPP-A, with a dissociation constant (Ko) of 61 pM (Figure S1F) and a half maximum inhibitory concentration (IC_50_) of 1.7 nM for mIGFBP-4 and 0.85 nM for mIGFBP-2 (Figure S1G). In contrast, it showed no binding activity to PAPP-A2, the closest homolog to PAPP-A (Figure S1F). To assess whether this antibody could block IGF signaling stimulated by the combination of IGF-1, IGFBP-4, and PAPP-A, we measured phosphorylation of AKT in HEK293T cells upon stimulation with IGF-1 in the presence of IGFBP-4 and PAPP-A. As expected, anti-PAPP-A blocked the downstream signaling of IGF-1 in HEK293 cells in a dose-dependent fashion, as measured by phosphorylation of AKT (Figure S1**H**, S1I). In vivo pharmacokinetic studies indicated that the antibody, formatted as an lgG1 isotype, has a half-life of 6.42 days in C57BL/6 mice (Figure S1J), allowing for weekly dosing. In summary, we have generated a PAPP-A neutralizing antibody with excellent potency, selectivity, and in vivo properties, allowing us to investigate the effects of pharmacologic PAPP-A inhibition.

### Anti-PAPP-A reduces collagen-associated extracellular matrix activity and IGF signaling in multiple tissues

PAPP-A KO mice have delayed age-related pathology across multiple tissues *(13)*, though the mechanism behind this extended longevity remains unknown. To establish a broad understanding of the effects of PAPP-A inhibition in adult wild-type animals, we treated male and female C57BL/6 mice with anti-PAPP-A or an isotype control antibody for 4 months, between 6 and 10 months of age, then analyzed a variety of tissues by RNA-seq. We used 5-6 mice/sex/group and analyzed 9 tissues from each animal (Figure 1A), for a total of 198 RNA-seq libraries.

**Figure 1:**
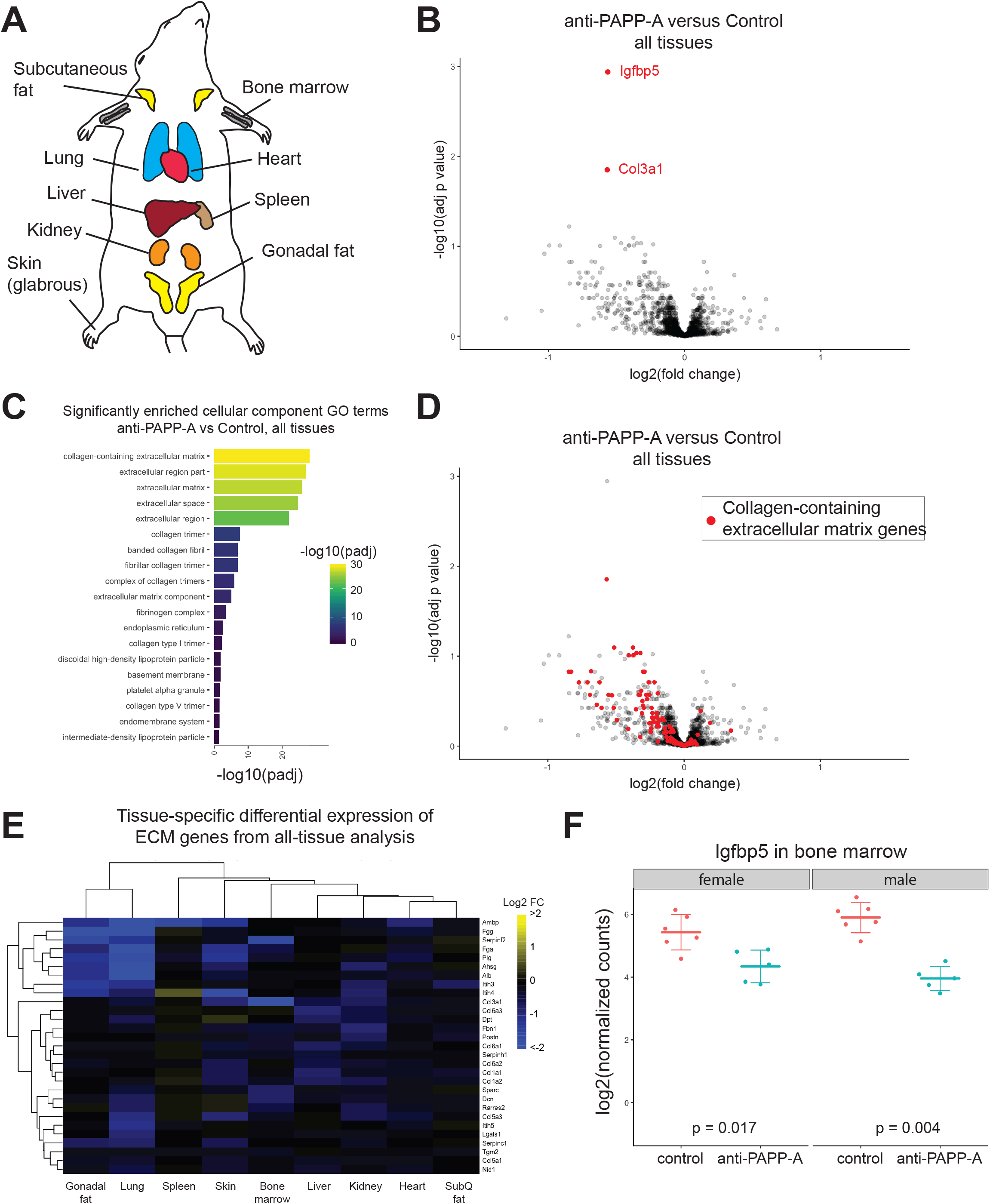
anti-PAPP-A reduces collagen-associated extracellular matrix activity and IGF signaling in multiple tissues. A. Diagram of the nine tissues (lung, liver, kidney, glabrous skin, gonadal fat, subcutaneous fat, spleen, heart, bone marrow) that were isolated and analyzed by RNA-seq from each animal, after 4 months of dosing. n = 6 male and 6 female for control antibody, 5 female and 5 male for anti-PAPP-A (198 total samples). B. Differential expression due to anti-PAPP-A across all tissues and both sexes, treating sex, tissue, and treatment as covariates. Red dots identify genes differentially expressed at FDR< 0.05. C. anti-PAPP-A affects collagen and ECM-related genes across multiple tissues. Cellular component GO terms significantly (FDR < 0.05) enriched in the list of genes differentially expressed by anti-PAPP-A at p < 0.01 in cross-tissue analysis (88 genes). D. Analysis of differential expression due to anti-PAPP-A across all tissues and both sexes. Red dots identify genes in the “collagen-containing extracellular matrix” gene set (GO:0062023). E. The tissue-specific log2 fold change for the 29 collagen-containing ECM genes that were affected by anti-PAPP-A across multiple tissues at p < 0.01. F. *lgfbp5* expression in whole bone marrow. Each dot represents the log2 (normalized counts) from RNA-seq data from an individual animal.

We were interested in identifying processes that might explain how PAPP-A affects multiple aspects of aging, so we performed differential expression analysis to identify genes that were affected by anti-PAPP-A similarly across all tissues, in both sexes. This stringent approach yielded only two genes that were differentially expressed at a 5% false discovery rate: *lgfbp5* and *Co/3a1*, both of which were decreased (Figure 1B). We therefore performed gene ontology (GO) analysis of all genes with a treatment p-value < 0.01 across both sexes and all tissues, consisting of 88 genes *(37)*. The results of this analysis demonstrated that the statistically significant reduction *Co/3a1* was only one component of a more coordinated downregulation of collagen-associated ECM-related genes: many of the significantly enriched (adj p < 0.05) GO terms in this set of 88 genes related to ECM and collagen biology, of which “collagen-containing extracellular matrix” was the most significant (adj p = 8.5e-29, 29 out of the 88 genes we examined were part of this gene set) (Figure 1C, S2A). This gene set consists of 334 genes, and visual inspection of the position of all these genes on the multi-tissue volcano plot showed a general decrease in gene expression across the entire set (Figure 1D), demonstrating that the GO term enrichment score above was not an artifact of any particular p-value cutoff. Further, this was not driven by any single tissue or small subset of tissues: when we examined the 29 affected genes from the “collagen-containing extracellular matrix” gene set in each tissue individually, we found that most tissues exhibited decreased expression of these genes, albeit with somewhat different expression patterns between tissues (Figure 1E). These results demonstrate that adulthood PAPP-A inhibition reduces collagen-associated ECM gene expression across multiple tissues. We hypothesized that this was via action on cells of mesenchymal origin, as mesenchymal cells are responsible for production of collagen and ECM in a variety of tissues *(38, 39)*.

It is notable that a non-collagen/ECM gene, *Igfbp5*, was the most significantly affected gene, as *Igfbp5* acts as a surrogate readout of IGF signaling: it is transcriptionally downregulated upon IGF inhibition and upregulated upon IGF stimulation in multiple cell types and tissues *(19, 20, 40–44)*. The observed decrease in *Igfbp5* expression due to anti-PAPP-A treatment demonstrates that IGF signaling was downregulated in multiple tissues, and indeed, when we examined each tissue and sex individually, *Igfbp5* trended towards decreased expression in multiple tissues in both males and females (Figure S2B). The most significant reduction of *Igfbp5* occurred in the bone marrow (Figure 1F), and this tissue also exhibited downregulation of multiple collagen and ECM genes, including *Co/3a1* (Figure 1E), which was the most significantly downregulated collagen gene across all tissues, as noted above. These results led us to focus on bone marrow as a relevant tissue to explore the mechanism and functional consequences of suppressed IGF signaling and collagen/ECM activity due to PAPP-A inhibition.

### Mesenchymal stromal cells are the primary source of PAPP-A in bone marrow and depend on PAPP-A activity for persistence

The bone marrow is the site of blood and immune cell production (hematopoiesis) and consists of both hematopoietic cells, produced by the rare but essential hematopoietic stem cells, and the stroma, which largely consists of mesenchymal and endothelial cell types (45, 46), including mesenchymal stromal cells (MSCs). MSCs are rare populations of non-hematopoietic, multipotent cells that are found across tissues in the stroma or perivascular niches and serve as progenitors of connective tissues (osteo-, adipo-, and condro-cyte lineages), as well as organizers of tissue niches (45, 47, 48). To characterize which bone marrow cells types were involved in IGF signaling and acutely responsive to PAPP-A inhibition, we treated 8 week old male C57BL/6 animals with a single dose of anti-PAPP-A or isotype control, isolated bone marrow from one week later, and then fractionated hematopoietic stem and progenitors (HSPCs) and mesenchymal stromal cells (MSCs). We further separated the HSPC population into hematopoietic stem cells (HSC), common lymphoid progenitors (CLP), granulocyte/macrophage progenitors (GMP), and megakaryocyte/erythrocyte progenitors (MEP), representing the major subclasses of myeloid and lymphoid differentiation (Figure 2A). Because most of these cell types are very rare (e.g. ~0.3% for MSCs, ≤0.05% for HSCs), we pooled the bone marrow from 5 mice per replicate, with 3-6 replicates per treatment group. We then assessed the transcriptome of each of these cell types via RNA-seq.

**Figure 2:**
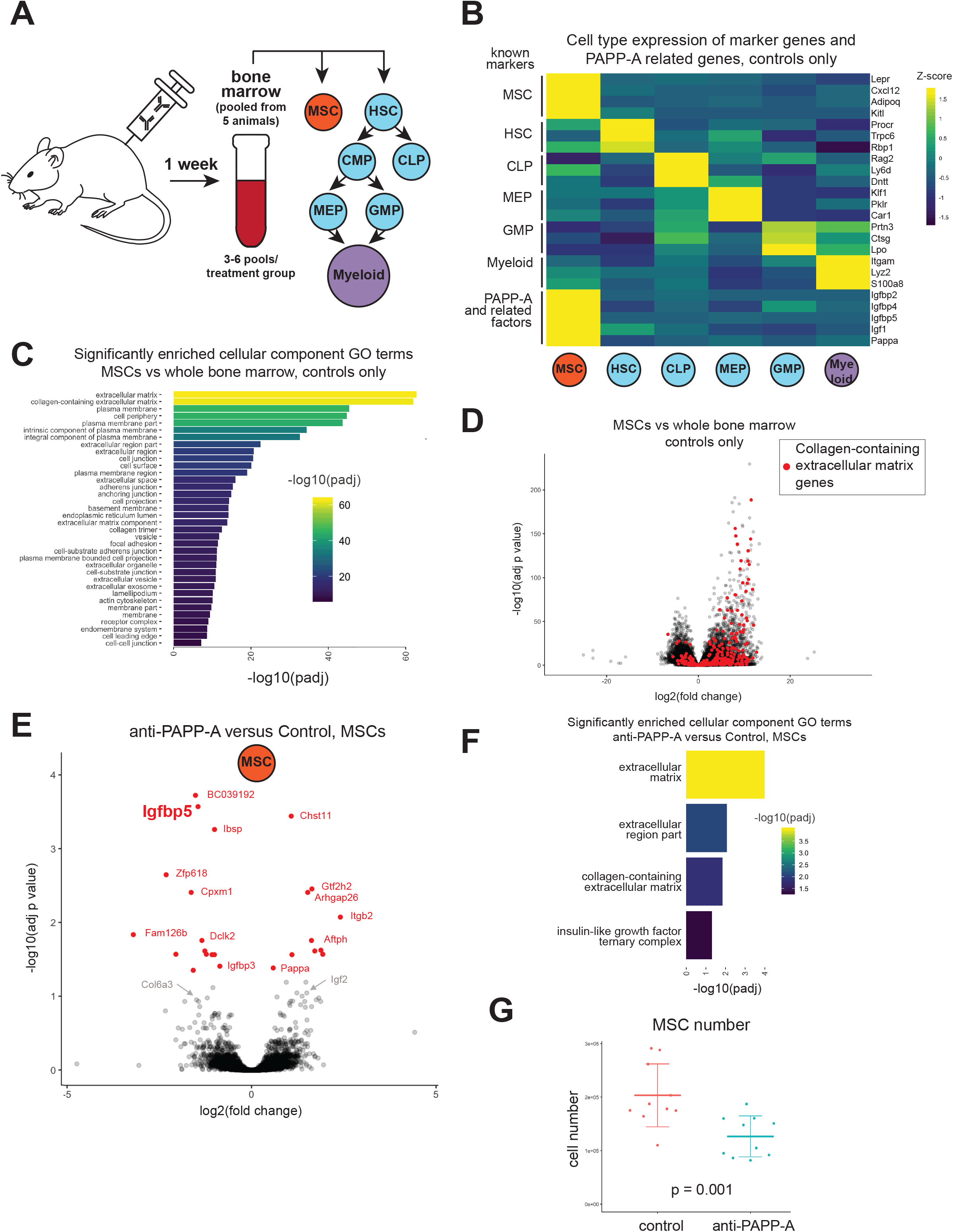
Mesenchymal stromal cells are the primary source of PAPP-A in bone marrow and depend on PAPP-A activity for persistence. A. Experimental design, indicating isolated cell types and their hierarchical relationship to one another. MSC= mesenchymal stromal cell (markers: CD31(-), CD45(-), Ter119(-), PDGFRa/CD140a(+), LepR(+)), HSC = hematopoietic stem cell (markers: Lin(-), c-Kit (+), Sca-1(+), CD48(-), CD150(+)), CMP = common myeloid progenitor (markers: Lin(-), Flk2(+), IL7Ra(+)), CLP = common lymphoid progenitor (markers: Lin(-), Flk2(+), IL7Ra(+)), MEP = megakaryocyte/erythrocyte progenitor (markers: Lin(-), c-Kit (+), Sca-1(-), FCgR(-), CD34(-)), GMP = granulocyte-macrophage progenitor (markers: Lin(-), c-**Kit** (+), Sca-1(-), FCgR(+), CD34(+)). B. Cell type specific expression of known marker genes and PAPP-A related genes. Control animals only. Colors represent z-scores of log2 normalized counts. C. MSCs express high levels of collagen and ECM-related genes. Cellular component GO terms enriched (FDR < 1e-7 for brevity) in the list of genes differentially expressed (FDR < 0.01, llog2FCI > 3) by MSCs versus whole bone marrow (2557 genes). Control animals only. D. Analysis of differential expression of MSCs versus whole bone marrow. Control animals only. Red dots identify genes in the “collagen-containing extracellular matrix” gene set (GO:0062023). E. Differential expression due to anti-PAPP-A in MSCs. Red dots identify genes differentially expressed at FDR < 0.05. F. anti-PAPP-A affects collagen/ECM related genes and IGF signaling in MSCs. Cellular component GO terms enriched (FDR< 0.05) in the list of genes differentially expressed by anti-PAPP-A at p < 0.005 in MSCs (105 genes). G. anti-PAPP-A reduces MSC cell number in bone marrow after 4 weeks of treatment (males only, n = 10/group).

Using control animals, we determined that this sorting paradigm effectively separated the cell types as assessed by known markers of each category, including the *Lepr, Cxc/12, Adipoq*, and *Kit/* markers of MSCs *(49–52)* (Figure 2B). Interestingly, when we examined the cell-type specific expression of *Pappa*, its substrates (*lgfbp2, lgfbp4, lgfbp5*), and its downstream effector, *Igf1*, we saw marked enrichment of expression of all these mRNAs in MSCs, as compared to HSPCs or myeloid cells (Figure 2B). Recent single-cell RNA-sequencing studies of murine bone marrow stroma have characterized non-hematopoietic cell populations in detail, and when we examined those datasets we found that, consistent with our results, *Pappa* was highly enriched in cells positive for the MSC markers *Lepr* and *Cxc/12 (49, 52)* (Figure S2C, S2D). Thus, we conclude that MSCs represent the primary source of PAPP-A in the bone marrow, as well as being major producers of other IGF signaling components.

Again using control animals, we examined the general pattern of gene expression in each cell type relative to whole bone marrow, focusing on MSCs because of their high PAPP-A expression. Thousands of genes were significantly differentially expressed in the sorted MSCs, and when we subjected the top genes (FDR< 0.01, llog2FCI > 3, 2557 genes) to GO term analysis, we saw a clear enrichment in GO terms related to extracellular matrix and collagen biology, of which “collagen-containing extracellular matrix” (the gene set most enriched by anti-PAPP-A in the cross-tissue analysis in Figure 1) was second most significant (adj p = 8.8e-47) (Figure 2C). Visual inspection of the 334 genes in this gene set on the MSC volcano plot indicated a general increase in gene expression across the entire set (Figure 2D). We conclude that MSCs are both the primary source of PAPP-A in bone marrow, as well as producers of the collagen and ECM genes downregulated by anti-PAPP-A. Of note, recent single-cell analyses have shown that MSC-derived cells, namely fibroblasts and osteolineage cells, are even greater producers of collagen/ECM in the bone marrow than MSCs themselves *(49, 52)*. The production of PAPP-A by MSCs and the production of collagen/ECM by MSCs and MSC-derived cells suggested that MSCs may be the primary mediators of the effect of anti-PAPP-A on collagen and extracellular matrix gene expression.

To test this hypothesis, we examined the response of each cell type to PAPP-A inhibition. HSPCs showed 22 genes significantly differentially expressed, none of which appeared to be related to IGF signaling or collagen/ECM production, and virtually all of which exhibited a less than a two-fold change (Figure S3A, Table S1). Analyzing each of the HSPC cell types individually (HSC, CLP, MEP, GMP) also yielded few significantly changed genes (Figure S3B, Table S2). Myeloid cells showed 11 genes whose expression changed significantly, again, without any apparent enrichment of IGF signaling or collagen/ECM production (Figure S3C, Table S2). In both cases, GO analysis yielded only general terms (e.g. nucleic acid binding) without strong enrichment. Notably, *Igfbp5* was not decreased in any of the HSPC or myeloid cell types (Figure S3D). In contrast, MSCs showed 24 genes whose expression changed, the second most significant of which was *Igfbp5* (Figure 2E). Interestingly, *Pappa* itself was slightly but significantly upregulated, and *Igf2* showed a trend toward upregulation, perhaps suggesting a partial compensatory response. We did not observe a striking downregulation of collagen/ECM genes, but when we submitted the top 105 genes affected by PAPP-A inhibition (p < 0.005) to GO term analysis, we saw significant enrichment for the same collagen and extracellular-matrix GO terms that we identified previously (Figure 2F), whereas no such enrichment was found for HPSC or myeloid cell types, irrespective of statistical cutoff. These data, particularly the notable reduction in *Igfbp5* after a single dose of anti-PAPP-A, support the hypothesis that MSCs are the proximal responders to anti-PAPP-A in the bone marrow.

Given the strong reduction in IGF signaling (measured by reduced *Igfbp5)* but relatively minor effect on collagen/ECM gene expression in sorted MSCs, we reasoned that the tissue-level reduction in collagen/ECM gene expression could have arisen through a reduction in total MSC number (and by extension, a reduction in the number of MSC-derived cells), which we would not have detected in the above analysis because we compared equal cell numbers/RNA content. To address this, we treated 8 week old male C57BL/6 animals with anti-PAPP-A or isotype control for 4 weeks and measured bone marrow MSC numbers. We observed a ~40% reduction in MSC number in both males and females, across two independent experiments (Figure 2G, S3E). To expand this finding, we also examined MSCs in PAPP-A KO mice, using immunophenotyping to quantify MSCs from the bone marrow of male and female PAPP-A KO mice at 3-6 months of age. We report frequency rather than absolute number, as PAPP-A KO mice are ~60% the size of WT mice. Similar to anti-PAPP-A treated animals, PAPP-A KO mice exhibited a significant reduction in MSC content (combined p = 0.035, Figure S3F), with no evidence of a sex-specific effect. Taken together, these results support a model in which anti-PAPP-A reduces IGF signaling in MSCs, which reduces their survival and/or proliferative ability and leads to a reduction in cell number over time. Because MSCs and their progeny are a major source of collagen and extracellular matrix factors, this leads to a reduction in collagen/ECM gene expression at the tissue level.

### PAPP-A inhibition reduces bone marrow myelopoiesis

Our data thus far suggested that PAPP-A inhibition altered MSC biology, but to test this at a functional level, we next examined downstream readouts of MSC activity. MSCs directly support hematopoiesis and HSCs in the bone marrow by providing cytokines and growth factors *(53, 54)*. Although we did not observe a direct effect of PAPP-A inhibition for 1 week on the transcriptome of HSPCs or myeloid cells, the important role of MSCs in forming the bone marrow niche led us to predict that a functionally relevant reduction in the number of MSC niche support cells would alter hematopoietic output. To test this, we treated young adult (2-4 month old), male C57BL/6 animals with weekly doses of anti-PAPP-A or isotype control and isolated bone marrow from the leg bones 4 weeks later (Figure 3A). We noticed that the anti-PAPP-A mice exhibited a trend toward reduced bone marrow cellularity (Figure S4A), and further examination of cell categories revealed that this loss was largely driven by a decrease in myeloid cells (CD11b+) (Figure 3B), which accounted for 71% of the lost cells (Figure 3C). Other major cell categories were either not reduced or reduced only slightly, including B cells (B220+), T cells (CD3+), HSPCs (Lin-, Ckit/CD117+), and other cell types that we did not specifically stain for (“Other”). Although we also noted a reduction in the non-hematopoietic MSCs, as mentioned above, these only made up ~0.3% of the overall bone marrow, and thus did not meaningfully contribute to overall decline in bone marrow cellularity. To test the robustness of this finding, we repeated the experiment with more animals and both sexes: n = 15 males and 15 females/group, again treated with anti-PAPP-A or isotype control for 4 weeks. Once again, bone marrow cellularity was reduced, significantly in females and non-significantly in males, though the average effect size was similar in both (Figure S4B). Again, this was largely driven by a reduction in myeloid cells in both sexes (Figure S4C). Of note, other cell types were also reduced in females, but myeloid cells remained a major contributor to the cellularity loss. Further suggesting this was primarily a myeloid effect, we assessed cellularity in the spleen (a major lymphoid organ that does not produce myeloid cells), and found no significant change in cellularity in these animals, either in total (Figure S4D) or when subdivided by cell category (Figure S4E).

**Figure 3:**
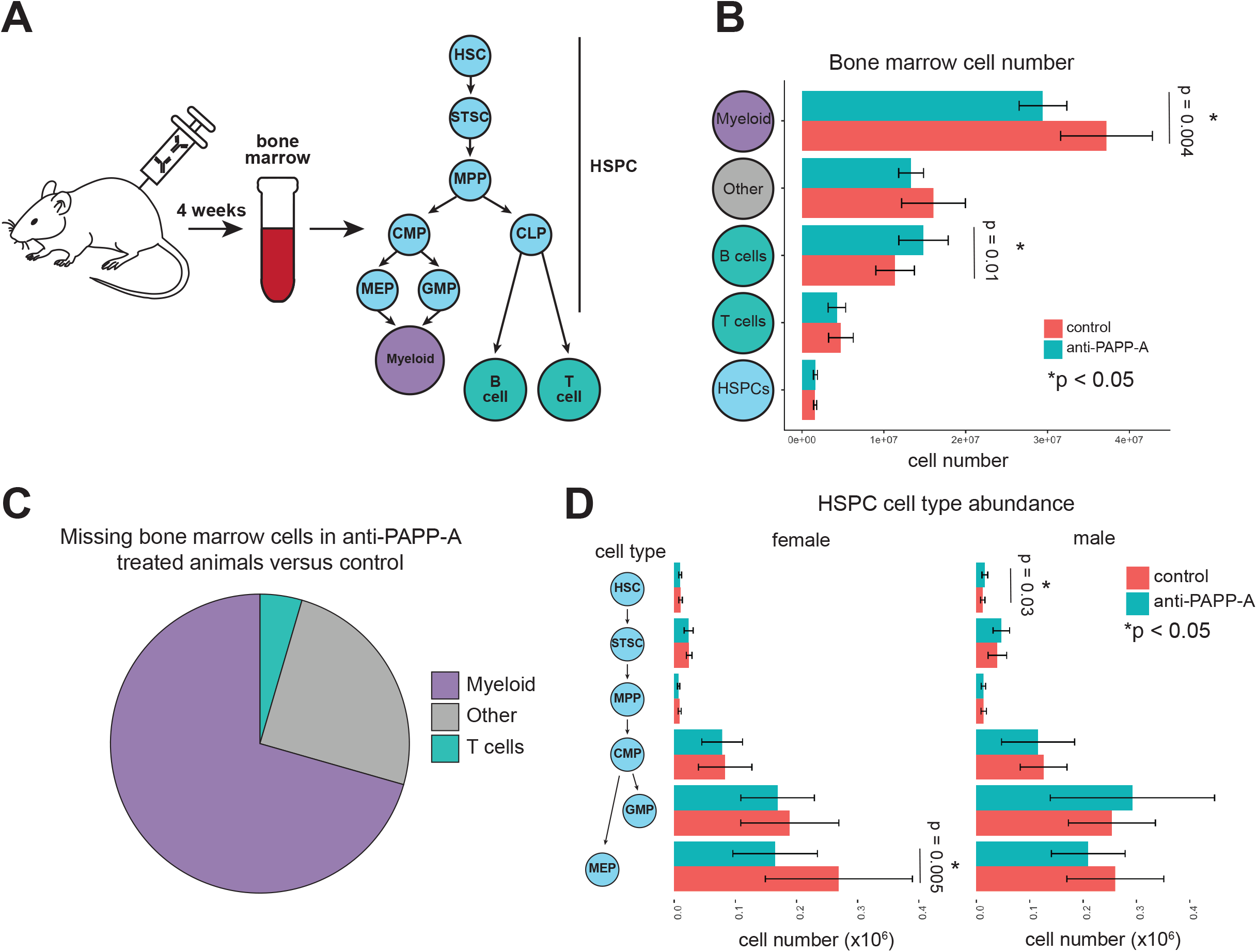
PAPP-A inhibition reduces bone marrow myelopoiesis. A. Experimental design, indicating isolated cell types and their hierarchical relationship to one another. MSC = mesenchymal stromal cell, HSC = hematopoietic stem cell, CMP = common myeloid progenitor, CLP = common lymphoid progenitor, MEP = megakaryocyte/erythrocyte progenitor, GMP = granulocyte-macrophage progenitor. B. PAPP-A inhibition reduces myeloid cell number in bone marrow. Abundance of each cell type in bone marrow after 4 weeks of treatment with anti-PAPP-A (males only, n = 10/group). C. Myeloid cells account for the majority of cells lost from the bone marrow after 4 weeks of PAPP-A inhibition. Same data as 3B, showing contribution of each cell category to missing cells in anti-PAPP-A group (males only, n=10/group). D. PAPP-A inhibition has minimal effect on HSPC abundance. Abundance of each cell type in bone marrow after 4 weeks of treatment with anti-PAPP-A (n = 15/sex/group).

To understand where in the myelopoiesis pathway this cell loss began, we examined hematopoietic stem and progenitor cells in the bone marrow in more detail. In both the original study (males only) and in the validation study (males and females), we saw no decrease in hematopoietic stem cells, short-term stem cells, or multipotent progenitors, which can differentiate into both myeloid and lymphoid lineages (Figure 3D, S4F). There was a significant reduction in megakaryocyte/erythrocyte progenitors (MEPs) in females and a trend in that direction in males, suggesting that the reduction in myelopoiesis may begin at this stage of differentiation, once myeloid/lymphoid commitment has occurred. Taken together, these results suggest that disruption of IGF signaling in MSCs and subsequent reduction in MSC number in the bone marrow niche primarily affects downstream myelopoiesis, with minimal effect on hematopoietic stem cells or other multipotent cell types, and minimal effect on lymphopoiesis.

### PAPP-A inhibition reduces osteogenesis via reduced osteoblast activity

In addition to providing hematopoietic niche support, bone marrow MSCs also give rise to most adult osteoblasts and are thus the major source of adult bone formation *(54)*. We reasoned that, if adulthood PAPP-A inhibition was functionally affecting MSCs, bone would be affected as well. Genetic modulation of PAPP-A has been shown to affect bone mass and geometry *(16, 55)*, but it was unclear whether short-term, adulthood inhibition of PAPP-A would result in bone effects. To assess this, we treated 18 male and 18 female C57BL/6 mice with anti-PAPP-A or isotype control for 4 months, starting at 6-8 months of age. We chose this age range to ensure that we were assessing adulthood bone homeostasis, rather than the bone growth that occurs during development, and thus focusing on the aspect of bone formation specifically affected by MSCs.

After 4 months of dosing, we isolated femurs and L5 vertebrae from these animals and analyzed them by micro computed tomography (µCT). PAPP-A inhibition resulted in a relatively mild, but highly significant reduction in multiple bone parameters in both sexes, both cortical and trabecular, in both the femur and the L5 vertebra (Figure S5A). The overall effect was sex-independent; although males trended towards a greater effect size than females, this may be a consequence of greater baseline bone retention in 6~8 month old males than females, allowing for a larger dynamic range. For cortical bone, the most consistent effect across both sexes was reduced cortical area (Figure 4A), and for trabecular bone, reduced trabecular thickness (Figure 4B) and associated parameters (e.g. bone surface/volume ratio, which increases as trabeculae become thinner).

**Figure 4:**
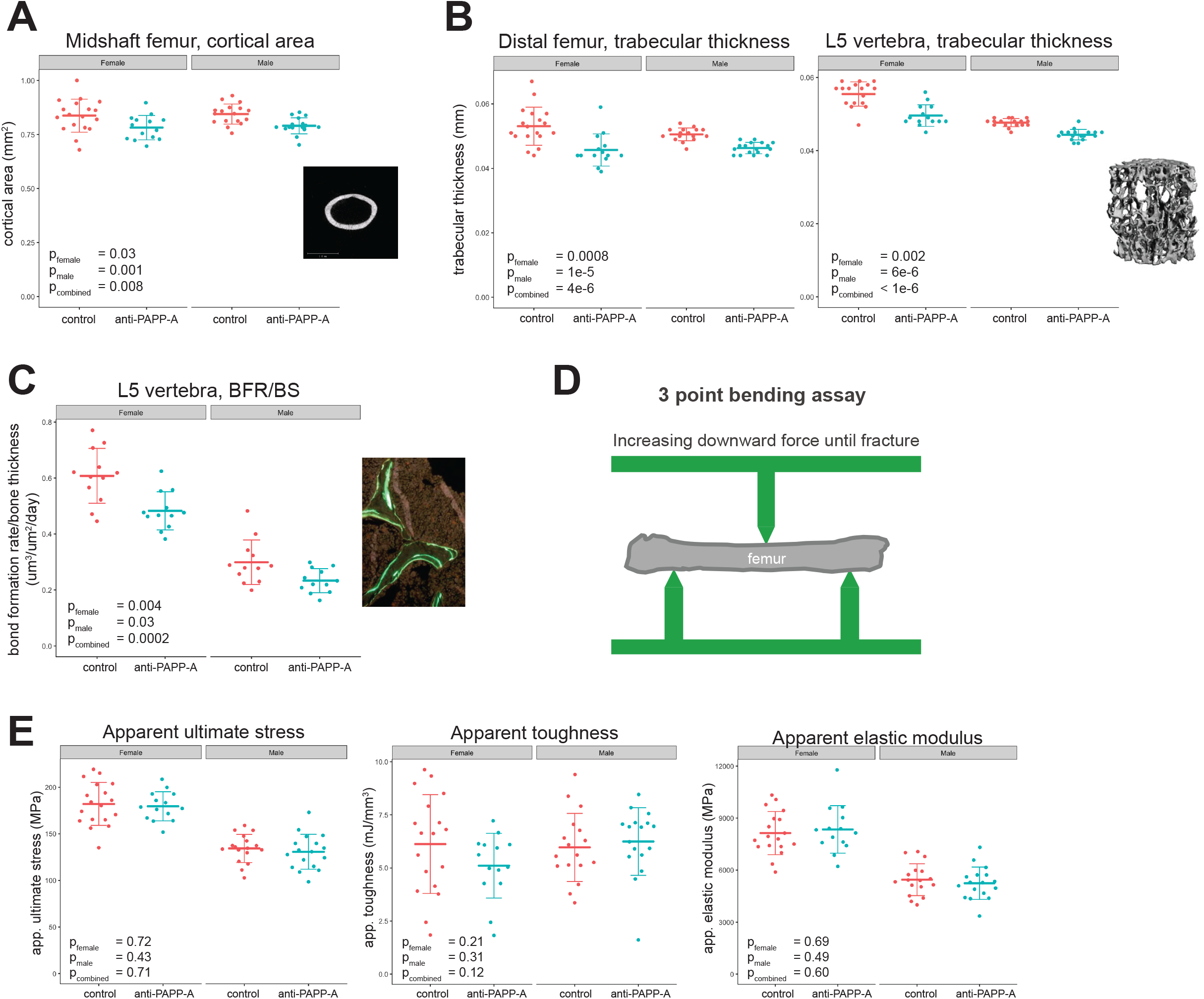
PAPP-A inhibition reduces osteogenesis via reduced osteoblast activity. A. Cortical area of midshaft femur, measured by uCT. Image shows a typical cortical scan, white area is quantified. B. Trabecular thickness in distal femur and L5 vertebra. Image is a 3D reconstruction of the trabecular bone in a typical L5 vertebral body. C. Bone formation rate (um^3^) per bone surface (um^2^) per day in L5 vertebra, assessed by dynamic histomorphometry. Image shows a typical histomorphometry image, BFR/BS is related to the distance between labels in double-labeled sections, with a larger distance indicating more bone formation. D. Schematic of three-point bending assay. Increasing downward force is applied until fracture, allowing for determination of multiple structural and apparent material properties. E. PAPP-A inhibition does not affect apparent bone material properties. Apparent material properties of femurs derived from three-point bend testing. App. ultimate stress is a geometry-normalized measure of the peak load applied to the bone; app. toughness is a geometry-normalized measure of the work the bone can absorb before fracture; app. elastic modulus is a geometry-normalized measure of the slope of the load-displacement curve prior to inelastic deformation. (n = 14-18/group for all experiments).

In both males and females, anti-PAPP-A resulted in a slight, but significant reduction in weight gain over time (Figure S5B). Bone mass and geometry are affected by body weight, so it was possible that the effect on bone µCT parameters was simply a result of reduced weight gain in the anti-PAPP-A animals. To account for this, we calculated q-values based on the residuals of each of the parameters accounting for individual animal body weight at takedown. For cortical bone, adjustment for weight effectively eliminated the effect, with no cortical parameters reaching q < 0.05, including cortical area (Figure S5C). In contrast, although adjustment for weight slightly reduced the statistical significance of anti-PAPP-A treatment on trabecular bone parameters, these effects remained highly statistically significant (Figure S5C), suggesting this was a direct effect on bone rather than a by-product of body mass.

The effect of PAPP-A inhibition on bone was consistent with the hypothesis that PAPP-A inhibition affected MSC-derived osteoblasts, but to more directly test this hypothesis, we performed dynamic histomorphometry on vertebrae and tibiae from these mice, which allows direct quantification of bone formation and resorption rates via injected fluorescent labels that mark bone formed over a known time window *(56)*. The most reliable histomorphometry data were obtained from the vertebra, as these retained measurable amounts of trabecular bone in both males and females. Males and females each exhibited either a significantly decreased or a trend toward decreased bone formation rate (Figure 4C), mineralizing surface (MS/BS), osteoblast number (N.Ob/B.Pm) and surface (Ob.S/BS) (Figure S5D). As there was no indication of a sex-specific effect, we performed combined-sex statistical analysis and the increased power resulted in a statistically significant decrease in bone formation (q = 0.003) and mineralizing surface (q = 0.046). The observations in the tibia are less reliable due to the almost complete disappearance of trabecular bone at this age, particularly in females, however bone formation rate (BFR/BS) and osteoblast numbers (N.Ob/B.Pm) and surface (Ob.S/BS) also trended toward a decrease in the anti-PAPP-A treated animals, with the latter measurements being significantly reduced in males (Figure S5D). In contrast to the reduced bone formation seen in trabecular bone, there was no effect on dynamic histomorphometry parameters in cortical bone (Figure S5D), consistent with the cortical effect being a mild, indirect consequence of lower body mass compared to the control group. Overall, the data show a consistent and coherent decrease in trabecular bone formation, driven by a decrease in both bone formation rate and osteoblast number, with no effect on bone resorption. Thus, the effects on trabecular bone microstructure seen above are most likely a consequence of reduced osteoblast number and activity, supporting the hypothesis that PAPP-A inhibition affects MSCs and their progeny (osteoblasts).

Having implicated PAPP-A in the biology of adult bone formation, we next asked whether this morphological change resulted in a functional weakening of bone, which could have implications for the safety of this potential therapeutic. We subjected the femurs examined by µCT above to mechanical testing via a 3 point bending assay (Figure 4D) to determine the material properties of the bone. Although, as noted above, the bones were slightly smaller (an effect that was highly correlated with weight), after adjustment for bone size, there were no differences between the treatment groups (Figure 4E), suggesting no deficit in the intrinsic properties of the bone material itself.

We also monitored circulating markers of bone formation/resorption over the course of the study. We measured Procollagen type 1 propeptides (P1NP) as a marker of bone formation and Tartrate-resistant acid phosphatase (TRACP) 5b as a marker of bone resorption. P1NP declined over the course of the study, consistent with previous reports and indicative of reduced bone formation with advancing age *(57)*. Females treated with anti-PAPP-A showed a slight, but significant, reduction in P1NP over time compared to controls, whereas in males, if anything, P1NP declined more slowly in anti-PAPP-A treated animals (Figure S5E). For TRACP5b, although there were differences between the groups that existed prior to treatment, there was no increased difference between the groups after treatment in either sex, suggesting no anti-PAPP-A effect (Figure S5F). To test another readout of bone resorption, we assayed circulating levels of C telopeptide of type 1 collagen (CTX-1) at the terminal timepoint and saw no effect in either sex (Figure S5G). In short, despite a decrease in bone formation and microstructure by highly sensitive readouts such as µCT and dynamic histomorphometry, PAPP-A inhibition did not have a substantial effect on circulating markers of bone formation or resorption, consistent with this being a relatively mild effect.

Taken together, these data demonstrate that adult PAPP-A inhibition reduces bone formation via a reduction in osteoblast number and activity. This effect is highly significant but does not seem to represent a pathological degree of bone loss, as indicated by no change to circulating markers of bone activity and no change in bone material properties. Along with the reduction in collagen/ECM gene expression and myelopoiesis shown earlier, these results demonstrate that the reduced IGF signaling in bone marrow MSCs, which leads to reduced MSC number and thereby bone forming cells, causes functional changes at the tissue level. This demonstrates that bone marrow MSCs require PAPP-A-mediated IGF signaling for normal activity in wild-type animals.

### PAPP-A is expressed by MSC-like cells in multiple tissues

MSCs have historically been isolated from the bone marrow, so bone marrow-derived MSCs represent the most-studied MSC subtype, but virtually all adult tissues contain a MSC-like population *(47, 48)*. These cells are often perivascular and serve as producers of collagen, extracellular matrix proteins, and adipogenic, fibrogenic, osteogenic, and chondrogenic cell types and there is increasing agreement that this broad cell category is adversely affected by aging and contributes to multiple types of pathology, particularly fibrosis and chronic inflammation *(39, 59–61)*. Because PAPP-A inhibition has beneficial effects on age-related pathology in multiple tissues *(13)*, we hypothesized that this arises through modulation of MSC-like cells in multiple tissues. The decrease in collagen-containing ECM genes in multiple tissues upon PAPP-A inhibition (from Figure 1) is certainly consistent with this hypothesis, and although testing the effect of PAPP-A inhibition on MSCs in multiple tissues is beyond the scope of this study, we took the first step by assessing whether PAPP-A was highly expressed in the MSCs of other tissues, as it is in bone marrow MSCs, using publicly available single cell RNA-seq (scRNA-seq) datasets.

The Tabula Muris is one of the broadest scRNA-seq datasets currently available, containing data for twenty mouse tissues *(62)*. In this dataset, mesenchymal cell populations from multiple tissues showed enrichment for *Pappa* gene expression, including stellate cells from the pancreas, which are thought to be a MSC-like population *(63)*, fibroblasts from the heart, and MSCs from trachea, adipose, lung, muscle, mammary gland (Figure S6A). Of note, the Tabula Muris bone marrow isolation procedure enriched for hematopoietic populations, therefore bone marrow MSCs are not part of this dataset. A similar multi-tissue scRNA-seq dataset was recently published using a different methodology than the Tabula Muris, and this dataset also showed strong enrichment of *Pappa* gene expression in mesenchymal cell populations from multiple tissues, including the bone marrow mesenchyme (CXCL12-high cells) and lung mesenchyme (mesenchymal alveolar niche cells and stromal cells) *(64)*.

We explored several other single-cell RNA-seq datasets, including one of vascular-associated cells in the mouse lung and brain *(65, 66)*. Consistent with perivascular MSC expression, this dataset showed *Pappa* gene expression enriched in “perivascular fibroblast-like cells” in both tissues (Figure S6B). These fibroblast-like cells were also strongly enriched for collagen and ECM expression, consistent with an MSC-like phenotype. We also examined a scRNA-seq dataset from human pancreas *(67)*. Similar to the mouse pancreas from the Tabula Muris, this dataset showed an enrichment of *PAPPA* gene expression in the MSC-like pancreatic stellate cells, which also expressed high levels of collagen and ECM genes (Figure S6C). These data demonstrate that multiple tissues in humans and mice harbor cells expressing relatively high levels of PAPP-A, and that these cells exhibit gene expression signatures, annotations, lineage, and anatomical location (perivascular) that are reminiscent of bone marrow MSCs.

## Discussion

In this study, we utilized a highly potent PAPP-A neutralizing antibody to determine that acute, adulthood PAPP-A inhibition reduces collagen/ECM gene expression in multiple tissues. Supporting a connection between PAPP-A and collagen/ECM production, it has recently been shown that overexpression of PAPP-A in the mouse mammary gland increased the amount, distribution, and orientation of collagen in post-partum breast tissue, and that injection of media from PAPP-A overexpressing cells into the mammary gland increased proliferation and collagen abundance *(68)*. Although we observed a decrease in collagen/ECM gene expression in multiple tissues, we chose to more deeply investigate the mechanism using bone marrow because bone marrow was the tissue most responsive to PAPP-A inhibition (as measured by downregulation of IGFBP5, a readout of decreased IGF signaling). We found that bone marrow MSCs are a major source of PAPP-A and depend on PAPP-A for activity: PAPP-A inhibition reduced IGF signaling in MSCs (measured by reduced *Igfbp5* expression), influenced collagen/ECM gene expression in MSCs themselves, and reduced the number of MSCs over a short period (4 weeks). To determine the functional consequences of PAPP-A inhibition, we assessed outputs of MSC activity - hematopoietic niche support and the generation of differentiated progeny (osteoblasts) - and found that anti-PAPP-A reduced these MSC functions. A similar mechanism is likely at work in other tissues, as we found that MSC-like cells from multiple other tissues expressed high levels of PAPP-A and collagen/ECM genes, and collagen/ECM gene expression was reduced by anti-PAPP-A in multiple tissues. We propose that PAPP-A is expressed by MSC-like cells in multiple tissues and potentiates IGF signaling in those MSCs, modulating MSC functions such as support of myeloid cell production, bone formation, and collagen/ECM deposition (Figure 5). These functions are important for normal tissue homeostasis, but with age, MSCs acquire altered gene expression signatures, becoming more myofibroblast-like, aberrantly expanding, and depositing more collagen and ECM *(28)*. Excessive collagen and ECM deposition are hallmarks of age-related pathology, and we speculate that reduced collagen/ECM production from MSCs is the mechanism by which PAPP-A inhibition extends longevity and delays aging in multiple tissues.

**Figure 5:**
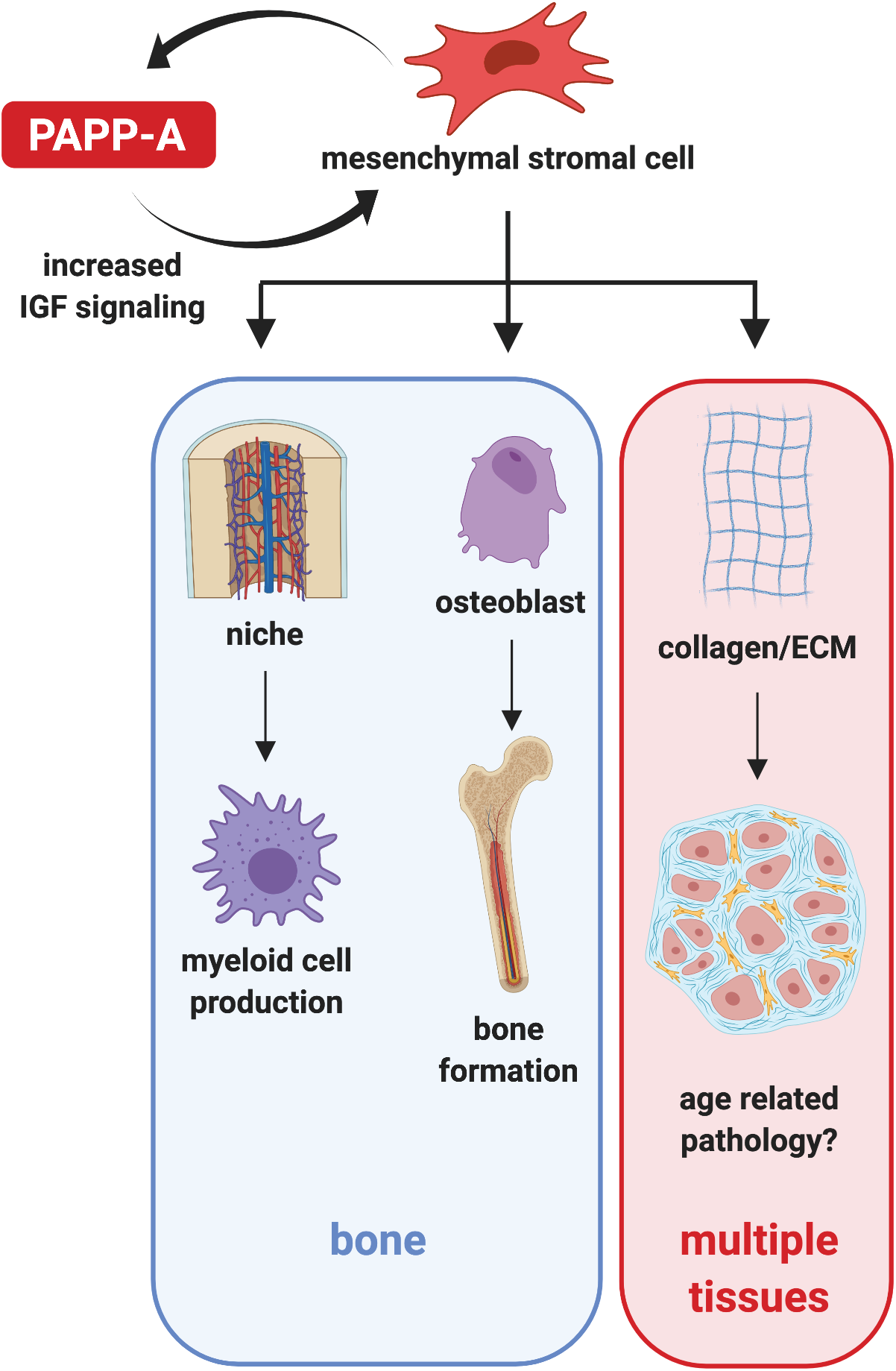
Model of PAPP-A mechanism of action. PAPP-A is expressed by MSCs and potentiates IGF signaling in those cells, increasing their activity and/or proliferation and promoting normal tissue function such as myeloid cell production, bone marrow, and collagen/ECM formation. With age, however, excessive collagen/ECM deposition from MSCs drives age-related pathology. This process is reduced by PAPP-A inhibition, potentially explaining the increased lifespan and healthspan in PAPP-A KO animals.

The production of PAPP-A by MSCs and their dependence on PAPP-A-mediated IGF signaling could explain many of the phenotypes of PAPP-A mutant mice. MSCs localize to the abluminal side of the vasculature, giving them a broad but relatively sparse tissue distribution *(47, 69)*. This patterning of MSCs may explain the ubiquitous yet very low levels of expression of PAPP-A found across tissues *(24)*. PAPP-A KO mice are proportional dwarves and their bones are correspondingly small *(16);* conversely, PAPP-A overexpression from the rat type I collagen promoter results in mice with increased bone thickness and bone size *(55)*. MSCs are the primary source of new osteoblasts in the adult, so it is reasonable to suppose that altered MSC activity may drive the skeletal and body size phenotypes of these PAPP-A mutants, though it is important to note that juvenile bone development is a very different process from adult bone homeostasis. PAPP-A KO mice are also reported to be resistant to fat accumulation on a high fat diet *(15, 16)*. Like osteoblasts, adipocytes are thought to derive from tissue-resident MSCs, and preadipocytes exhibit many similarities to bone marrow MSCs *(70);* thus, reduced adiposity due to PAPP-A inhibition may be a consequence of reduced number and/or size of MSC-derived adipocytes. PAPP-A KO mice are also resistant to atherosclerotic plaque progression and neointima formation without modulation of lipid levels *(18–20)*. There is evidence to suggest that plaque progression is potentiated by inflammation-driven activation of a pro-ECM phenotype in perivascular MSC-like cells *(71)*. It is plausible that PAPP-A inhibition delays this process by reducing the activation or expansion of these cells. Alternatively, it is possible that the effects of PAPP-A inhibition on aging and age-related disease derive primarily from the effects on myelopoiesis, i.e., the decrease in myeloid cells in the bone marrow upon PAPP-A inhibition. This hypothesis is attractive because aberrant innate immune activation and age-related myeloid skew are thought to drive multiple aspects of age-related decline, however, although we observed a clear decrease in myeloid populations within the bone marrow, we did not consistently observe equivalent changes in circulating myeloid populations (perhaps suggesting a compensatory mechanism at work), therefore we do not currently favor this hypothesis.

The association of PAPP-A expression with age-related pathologies in humans may be linked to its MSC origin and role in MSC biology. MSCs provide structural support via the production of ECM components like collagen and proteoglycans, but when ECM is laid down in excess or not removed properly once injuries are resolved, fibrosis can occur. Aberrant MSC activation has been implicated in fibrosis in multiple organs, including kidney, heart, lung, liver, skin, and bone marrow *(28)*. The expansion of these cells may explain the increase in *PAPPA* and *IGFBP* gene expression seen in human fibrosis, including pulmonary and hepatic fibrosis *(72, 73)*. Circulating levels of PAPP-A are elevated in patients with acute coronary syndrome and are predictive of poor outcome *(74–76);* the source of this circulating PAPP-A remains unknown, but may come from inflammation-activated perivascular MSCs, smooth muscle cells, or myofibroblasts, as PAPP-A has been shown to increase in response to cytokine stimulation in multiple mesenchymal cell types *(11, 26, 27)*. Lastly, high *PAPPA* gene expression is associated with poor outcome in breast cancer patients, and patients with high *PAPPA* (as well as high *SNA/1* and *COL1A*1) show a differential gene expression signature in which the top upregulated pathways are extracellular matrix organization, collagen formation, and epithelial-to-mesenchymal transition *(68)*. Suggesting the association is causal, PAPP-A overexpression in human breast cancer cells in vitro increased proliferation and collagen gene expression *(68)*.

The results of this study open new avenues of research around PAPP-A and IGF signaling. Now that we know PAPP-A is expressed in MSCs, it will be important to determine if PAPP-A is expressed by all MSCs or certain subsets. The expanding quantity of available single-cell RNA-seq data will help address this question, and focusing on MSC-specific transcription factors and transcriptional profiles may help answer the long-standing question of how PAPP-A is transcriptionally regulated. It is also important to determine whether all MSCs, or only certain subsets, require PAPP-A for persistence and activity. This will be best addressed via in vitro studies and/or a flexed allele of PAPP-A and cell-type specific Cre lines. Further, the exact mechanism by which MSCs are lost from the bone marrow upon PAPP-A inhibition remains unclear. It is possible that MSCs require IGF signaling for survival, and that the reduction in MSC number upon PAPP-A inhibition is a consequence of induced apoptosis. Increased IGF signaling has been reported to promote bone marrow MSC survival *(77)*, so it is reasonable to propose that reduced IGF signaling may reduce survival. Alternative interpretations are that a loss of PAPP-A-mediated IGF signaling reduces MSC proliferation, causes expulsion from the bone marrow, or induces differentiation (although the reduction in MSC-derived osteoblasts upon PAPP-A inhibition suggests this last option is unlikely). Future experiments will determine which of these mechanisms is primarily responsible. The field would also benefit from a more thorough investigation of the causal chain between PAPP-A inhibition, alteration to MSC number and activity, and functional outcomes. For example, it is possible that the effects of PAPP-A inhibition on hematopoiesis and bone formation are direct, independent of MSCs. We do not believe this to be the case, as we saw no evidence of altered IGF signaling in hematopoietic populations, and it is known that MSCs are required for normal organization of the bone marrow perivascular niche microenvironment, which supports hematopoiesis, and that MSCs themselves serve as a key source of HSC trophic factors and directly support HSCs *(53, 54, 78)*. Similarly, osteoblasts are known to arise from MSCs, and MSCs are the major source of adult bone *(54)*. However, we cannot rule out the possibility of an MSC-independent effect; this could be more thoroughly addressed via scRNA-seq of the entire bone marrow upon PAPP-A inhibition and phenotyping of mouse models with deletion of PAPP-A or IGF-1R only in specific cell types. To the extent that the effects of PAPP-A inhibition are indeed caused by MSC modulation, it remains unclear whether the relevant effect is a change in MSC number, a change in MSC activity state, or both. Studying changes to MSCs in multiple tissues during aging in the context of PAPP-A inhibition would greatly expand our understanding of this biology.

In summary, the established phenotypes of PAPP-A deficient mice coupled with the new knowledge described here, that PAPP-A modulates MSC activity, hint that there are many therapeutic avenues for PAPP-A inhibitors. From a safety perspective, the role of PAPP-A as a fine-tuner of IGF signaling is beneficial: both IGF signaling and MSCs play fundamental roles in normal tissue homeostasis, such that complete inhibition/ablation would likely be deleterious. PAPP-A neither eliminates IGF signaling nor completely ablates MSCs; rather, it tunes their activity such that normal functions remain (as evidenced by the health of PAPP-A KO mice), but may prevent aberrant activation in the context of injury and aging. Reduced IGF signaling has been a compelling longevity target for decades; PAPP-A inhibition seems to provide a safe and effective path toward that goal, and this study represents a major step forward in our understanding of the relevant cellular mechanisms.

## Methods

### Study design

Previous studies have implicated PAPP-A inhibition in the extension of lifespan and healthspan, but the mechanism behind this effect is unknown, as is the effect of shorter-term PAPP-A inhibition. The main goal of our study was to evaluate the cellular and molecular effects of a PAPP-A neutralizing antibody in vivo. Using this antibody, we demonstrated that PAPP-A inhibition reduced collagen and ECM gene expression in multiple tissues. We further investigated the bone marrow and found that PAPP-A inhibition reduced mesenchymal stromal cell number and activity, as well as altering tissue-level bone marrow phenotypes, including hematopoietic output and bone parameters. Sample sizes were calculated by power analysis from pilot studies and/or by our experience measuring the pre-specified endpoints. Mice were randomized to treatment group by body weight. For most experiments, investigators were blinded to which intervention each animal received, and measurements (e.g., bone parameters, ELISAs, FACS-based cell counts) were blinded until statistical analysis. Endpoints were chosen on the basis of our expertise and were prospectively selected. In cases where multiple endpoints were assessed, FDR correction has been applied. No outliers were excluded from analysis.

### In vitro PAPP-A enzymatic cleavage of IGF binding proteins

Human and murine PAPP-A proteins with C-terminal C-myc and Flag tags were expressed by stably transduced HEK293 cell line and purified by heparin column chromatography. Human and murine IGFBP-2, IGFBP-4 and IGFBP-5 proteins were produced recombinantly by transient expression in HEK293 cells with either N-terminal (for human proteins) or C-terminal (for murine proteins) 6His tag and purified by Ni-Sepharose column chromatography. For enzymatic cleavage reaction IGFBP-2 and IGFBP-4 proteins were pre-incubated with IGF1 of appropriate species (R&D Systems, 291-G1-200 for human and 791-MG-050 for mouse) for 30 min at 37°C. IGFBP-2/IGF1, IGFBP-4/IGF1 or IGFBP-5 proteins were then mixed with various concentrations of PAPP-A and incubated for 2-4 hours at 37°C. Final concentration in cleavage reactions were 90 nM for IGFBPs and 850 nM for IGF1. Proteins were then resolved by capillary electrophoresis on Wes instrument (ProteinSimple) using capillary cartridge kit (ProteinSimple, cat. # SM-W002-1), probed with **THE** HisTag antibody (GeneScript, cat. # A00186) and visualized with anti-mouse detection module (ProteinSimple, cat.# DM-002). To evaluate neutralizing potency of anti-PAPP-A antibody, PAPP-A protein was pre-incubated with various concentrations of anti-PAPP-A antibody prior to adding to IGF1/IGFBP-4 mix. PAPP-A concentration is these assays was fixed to 0.4 nM for IGFBP-4, 3.5 nM for IGFBP-2 and 0.08 nM for IGFBP-5 cleavage.

### Cellular pAKT assay

IGFBP-4 protein was mixed with IGF1 and incubated for 30 min at 37°C. PAPP-A protein was added to IGFBP-4/IGF1 mix and incubated for 4-5 hours at 37°C. Final concentrations in the reaction were 90 nM for IGFBP-4, 15 nM for IGF1 and 0.6 nM for PAPP-A. HEK293 cells were plated in EMEM media without serum and allowed to adhere overnight. IGF1/IGFBP-4/PAPP-A mix was added to cells at 1:30 final dilution and incubated for 20 min at 37°C. Cells were lysed in MSD Tris Lysis buffer and analyzed by Phospho(Ser473)/Total Akt Whole Cell Lysate kit (MSD, K15100D) according to manufacturer’s protocol. To evaluate neutralizing potency of anti-PAPP-A antibody, PAPP-A protein was pre-incubated with various concentrations of anti-PAPP-A antibody prior to adding to IGF1/IGFBP-4 mix.

### Gene expression analysis of cell lines

IMR-90 human lung fibroblasts (ATCC CCL-186), Hela human cervical epithelial cells (ATCC CRM-CCL-2), and 293T human embryonic kidney epithelial cells (ATCC CRL-3216) cell lines were procured from ATCC and cultured as recommended by ATCC. Hela cells were plated in 10% serum containing media, allowed to adhere overnight, then the next day were switched to serum free medium for 24hrs. Cells were then treated with one or a combination of the following recombinant proteins in serum free media [recombinant human IGF-1 (Gibco#PHG0071), recombinant human IGFBP-4 (R&D #804-GB), and recombinant human PAPP-A (R&D #2487-ZN (aa 82-1627 full length)] at the ratio of 5nM: 20nM: 2nM (IGF-I: IGFBP-4: PAPP-A) for 60 minutes. Cells were then washed and RNA was collected using TRIzol.

### Antibody production and validation

Anti-PAPP-A antibody with mouse IgG1 isotype was produced by transient transfection of HEK293 cells and purified by Protein A chromatography. The protein was estimated to be 98.9 % monomer by SEC chromatography and had Endotoxin of less than 0.27 EU/mg. Binding kinetics of anti-PAPP-A antibody to both human and mouse PAPP-A was evaluated by SPR with antibody capture on chip CM5 and PAPP-A concentrations ranging from 0.78 to 100 nM. Pharmacokinetic profile of anti-PAPP-A antibody was evaluated by single dose IV injection of 5 mg/kg in C57BL/6 male mice. Antibody blood concentration was measured by direct ELISA with PAPP-A protein.

### Mice

Male and female C57BL/6 wild type (WT) mice were purchased from The Jackson Laboratory (strain #000664) at approximately 3-6 months of age. Once at Calico and AbbVie, mice were housed 3-5 mice to a cage, and underwent a minimum 3-day acclimation period prior to placement on study. Mice were maintained on a 12:12 hour light:dark cycle at 22°C with environmental enrichment (e.g., nesting material and/or huts) with access to water and food ad libitum. All studies were performed in adherence to the NIH Guide for the Care and Use of Laboratory Animals.

PAPP-A knock-in reporter mice were generated by The Jackson Laboratory using standard CRISPR/Cas9 methods. Briefly, a P2A-HA-Luciferase-EGFP-bGHpA cassette was inserted inframe into exon 2 of *Pappa* (NC_000070.6). These mice report PAPP-A expression while disrupting expression of endogenous *Pappa*, thus heterozygous mice express one copy of PAPP-A and one copy of the reporter while homozygous reporter mice express two copies of the reporter and zero copies of PAPP-A and are thus essentially PAPP-A knockouts (KO).

WT mice were treated with control (anti-TeTx, lgG1) or anti-PAPPA antibody (lgG1) at 10 mg/kg via IP injection once a week for the duration of each study.

### Preparation of tissues and isolation of cells by flow cytometry

Hematopoietic stem and progenitor cells (HSPCs) were analysed and/or sorted as described *(79, 80)*. To obtain bone marrow cells, mice were sacrificed via CO_2_ asphyxiation followed by cervical dislocation. For bone marrow analysis, mice were analyzed one at a time and only leg bones were used (2 femur and 2 tibia per mouse). For bone marrow sorting, groups of 5 mice were pooled and all major bones were used (2 femur, 2 tibia, 2 ilium, 2 humerus, 2 radius, 2 ulna per mouse). Bone marrow cells were harvested in staining media (SM) (PBS without Ca^2^+ or Mg^2^+ supplemented with 2% fetal bovine serum (FBS)) by gently crushing bones with a mortar and pestle and filtering through 40um cell strainers. Erythrocytes were lysed in ACK (150mM NH4Cl/10mM KHCO3) lysis buffer and then remaining cells were washed in SM. Bone marrow cells were counted using a Vi-Cell and then aliquoted for staining for FACS. Only for sorting, bone marrow cells were purified on a Ficoll gradient (Histopaque 1119, Sigma-Aldrich, #11191-100ml), to remove bone dust and dying cells. Cells were then enriched for HSPCs by using c-Kit microbeads (Miltenyi Biotec, #130-091-224) and MACS Separation LS Columns (Miltenyi Biotec, #130-042-401).

Blood was collected via retro-orbital bleed or via terminal cardiac puncture and collected in EDTA-coated tubes (Becton Dickinson). Erythrocytes were lysed in ACK lysis buffer and then remaining cells were washed in SM. Blood cells were then stained for FACS analysis.

Spleens were dissociated by gently pressing spleen through a 40um cell strainer with the plunger of a 3ml syringe and washing with SM. Erythrocytes were lysed in ACK lysis buffer and then remaining cells were washed in SM. Spleen cells were counted using a Vi-Cell and then aliquoted for staining for FACS analysis.

To analyze or sort HSPCs, cells were stained with the following antibodies: biotinylated lineage markers [Mac-1 (CD11b), Gr-1 (Ly-6G/C), Ter119 (Ly76), CD3, CD4, CD5, CD8a (Ly-2), and B220 (CD45R) (Biolegend)] and streptavidin conjugated BUV-395, IL?Ra (CD127) BUV-737, FCgR BV-711, CD48 BV-605, Flk2 (CD135) BV-421, Sca-1 (Ly-6A/E) PE-Cy?, CD150 PE-594, CD34 PE, Ckit APC-Cy7. To analyze or sort mature immune cells, cells were stained with the following antibodies: CD11b BUV-737, B220 BUV-496, F4/80 BV-650, Gr1 BV-421, CD3 PE. To analyze or sort MSCs, bone marrow cells were stained with the following antibodies: CD45 BUV-395, CD31 BUV-395, Ter119 BUV-395, CD140a APC, Biotinylated LepR and Streptavidin PE.

MSCs were isolated as described *(49, 66, 81)*. Briefly, bones were crushed using a mortar and pestle in SM. Both marrow and bone fragments were then dissociated in 1mg/ml Stemxyme (Worthington, #LS004106), 1mg/ml Dispase II (ThermoFisher Scientific, #17105041), and DNAsel (200U/ml, Roche, #10104159001) in SM at 37°C for 25 minutes shaking. Cells were then filtered sequentially through 110um and 70um cell strainers, washed and then erythrocytes were lysed using ACK-lysis buffer. Cells were then labeled with Dyna-bead conjugated antibodies that had been prepared the night before. To prepare antibody-Dynabead conjugates, 200ul Sheep Anti-Rat IgG Dynabeads (Thermofisher Scientific, #11035) per mouse were incubated overnight at 4°C shaking with Rat-anti mouse antibodies against CD45, CD31, Ter119, B220, CD3, CD4, CD5, CD8, Gr1, and CD11b. The next day, just before adding prepared cells, tubes were put on Dyna5 magnet (ThermoFisher Scientific, #12303D) and the supernatant poured off. Then prepared cells were added to the antibody-Dynabead conjugates (no longer on the magnet) and incubated at 4°C shaking for 45 minutes. Tubes were then put back on the Dyna5 magnet and the negative fraction was collected. The cells from the negative fraction were then stained with CD45-BUV395, CD31-BUV395, Ter119-BUV395, CD140a-APC, Biotinylated LepR and Streptavidin PE, and MSCs were sorted.

Flow cytometry analysis was performed on a BD Fortessa and flow cytometry cell sorting was performed on a BD FACS ARIA. All data analysis was done using BD Diva and Treestar FlowJo software programs.

Detailed description of antibodies used for flow cytometry:

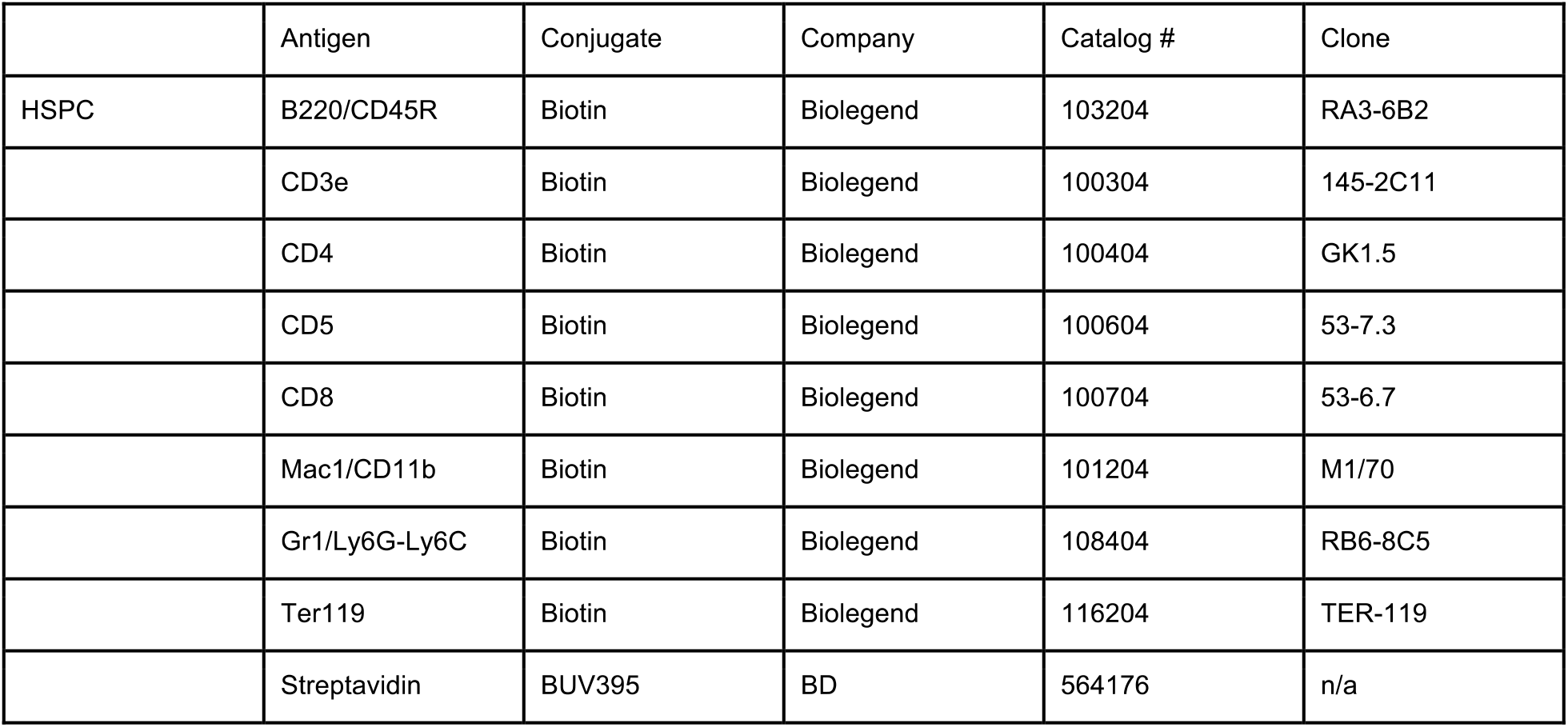
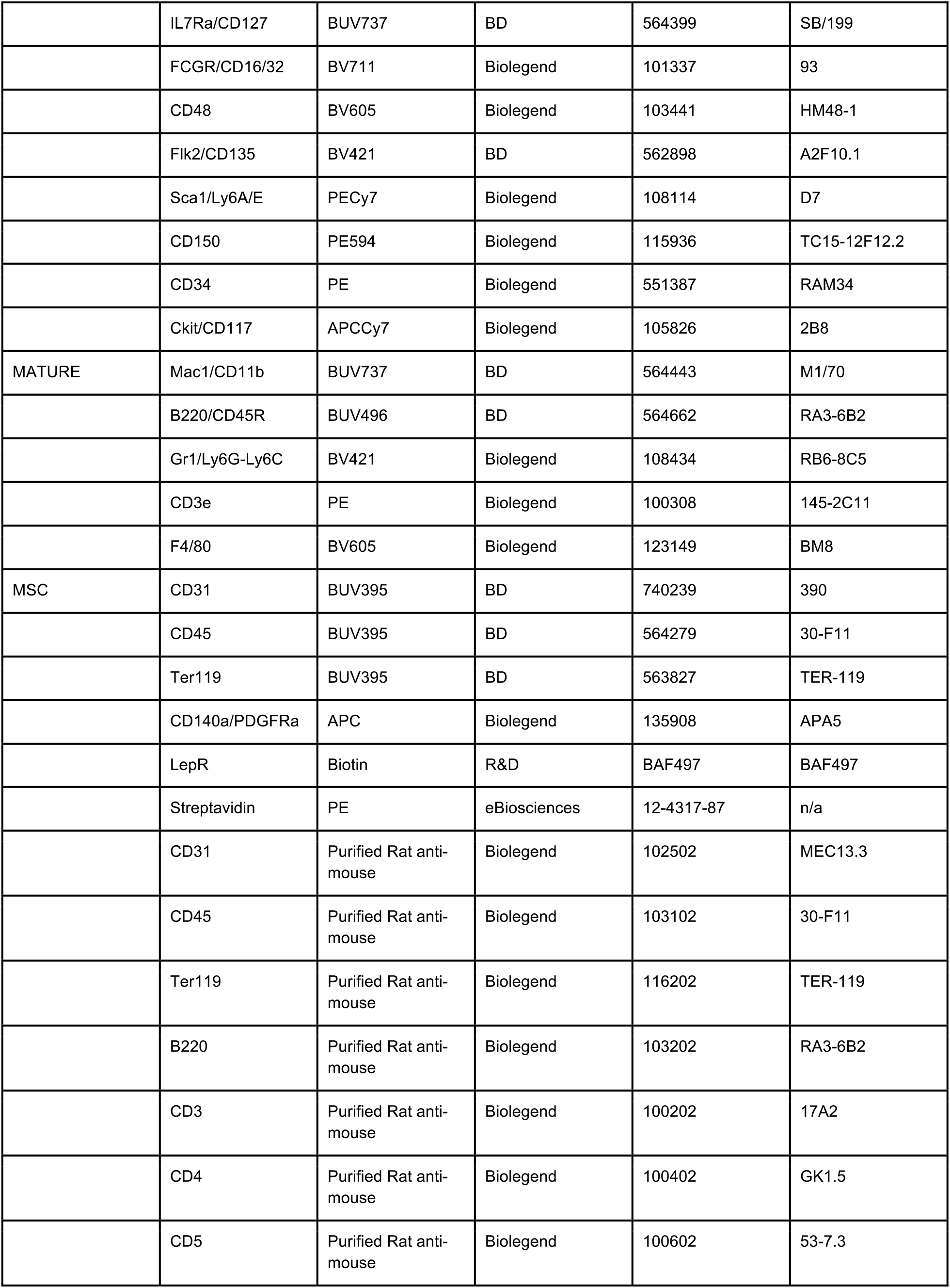
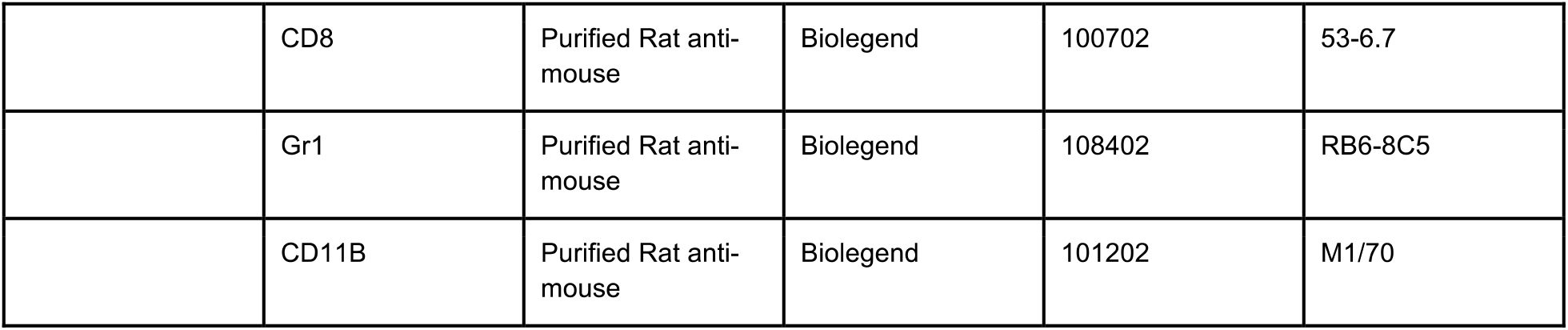

### Gene expression

RNA was extracted from whole tissues using the Qiagen RNeasy 96 Universal Tissue Kit (Qiagen, 74881) and was digested with DNAsel before preparation for RNA-seq. RNA was extracted from low numbers of sorted populations using TRlzol reagent (lnvitrogen) and Linear Acrylamide (ThermoFisher, #AM9520) was used to aid precipitation and pellet visualization, RNA was then digested with DNaseI before preparation for RNA-seq.

Sequencing libraries were prepared as directed using either TruSeq® Stranded Total RNA Library Prep Human/Mouse/Rat (Illumina) or NEBNext® Ultra™ II Directional RNA Library Prep Kit (NEB). Samples were amplified for 10-13 cycles and sequencing on a HiSeq 4000 (Illumina). Sequencing quality control was performed using FastQC (v0.11.5). Transcript expression was then quantified using Salmon (v0.9.1) *(82)* in pseudo-alignment mode, without adapter trimming, producing per-gene count estimates, using the Ensembl mm10 transcriptome. Normalized count and differential expression analysis were performed in R using DESeq2 (v1.22.2) *(83)*, with a pseudo-count of 0.5 added to all normalized counts. For the cross-tissue analysis in Figure 1, treatment effect was extracted from a model incorporating sex and tissue as covariates (value ~ sex + treatment + tissue).

### Bone analyses

#### µCT

High resolution micro-computed tomographic imaging (µCT40, Scanco Medical AG, Brüttisellen, Switzerland) was used to assess microarchitecture and mineral density in the femur and fifth lumber (LS) vertebra of each mouse. In the femur, trabecular bone microarchitecture and cortical bone morphology were measured in the distal femoral metaphysis and mid-diaphysis, respectively. Trabecular bone was analyzed in the L5 vertebral body. Scans were acquired using a 10 µm^3^ isotropic voxel size, 70 kVP peak x-ray tube potential, 114 mAs tube current, 200 ms integration time, and were subjected to Gaussian filtration and segmentation. Image acquisition and analysis protocols adhered to the JBMR guidelines for the assessment of rodent bones by µCT *(84)*. Trabecular bone was analyzed in the distal femur and L5 vertebral body. In the distal femur, transverse µCT slices were evaluated in a region of interest (ROI) beginning 200 µm superior to the distal growth plate and extending proximally 1500 µm (150 slices). Trabecular bone in the L5 vertebral body was analyzed in ROI that extended from 100 µm inferior the cranial endplate to 100 µm superior to the caudal end-plate. The trabecular bone region was identified by semi-manually contouring the trabecular bone in the ROI with the assistance of an auto-thresholding software algorithm. Images were thresholded using an adaptive-iterative algorithm. The average adaptive-iterative threshold of the Female Control mice (375 mgHA/cm^3^) was then used as the threshold to segment bone from soft tissue in all samples. Morphometric variables were computed from the binarized images using direct, 3D techniques that do not rely on any prior assumptions about the underlying structure. For trabecular bone regions, we assessed the bone volume fraction (Tb.BV/TV, %), trabecular bone mineral density (Tb.BMD, mgHA/cm^3^), trabecular specific bone surface (Tb.BS/BV, mm^2^/mm^3^), trabecular thickness (Tb.Th, mm), trabecular number (Tb.N, mm-1), trabecular separation (Tb.Sp, mm), connectivity density (ConnD, 1/mm^3^), and structure model index (SMI), a parameter that describes the plate-vs-rodlike nature of the architecture. Higher values of SMI indicate rod-like structures, whereas lower values characterize plate-like ones. See below for a description of each of the parameters:

Trabecular bone parameters

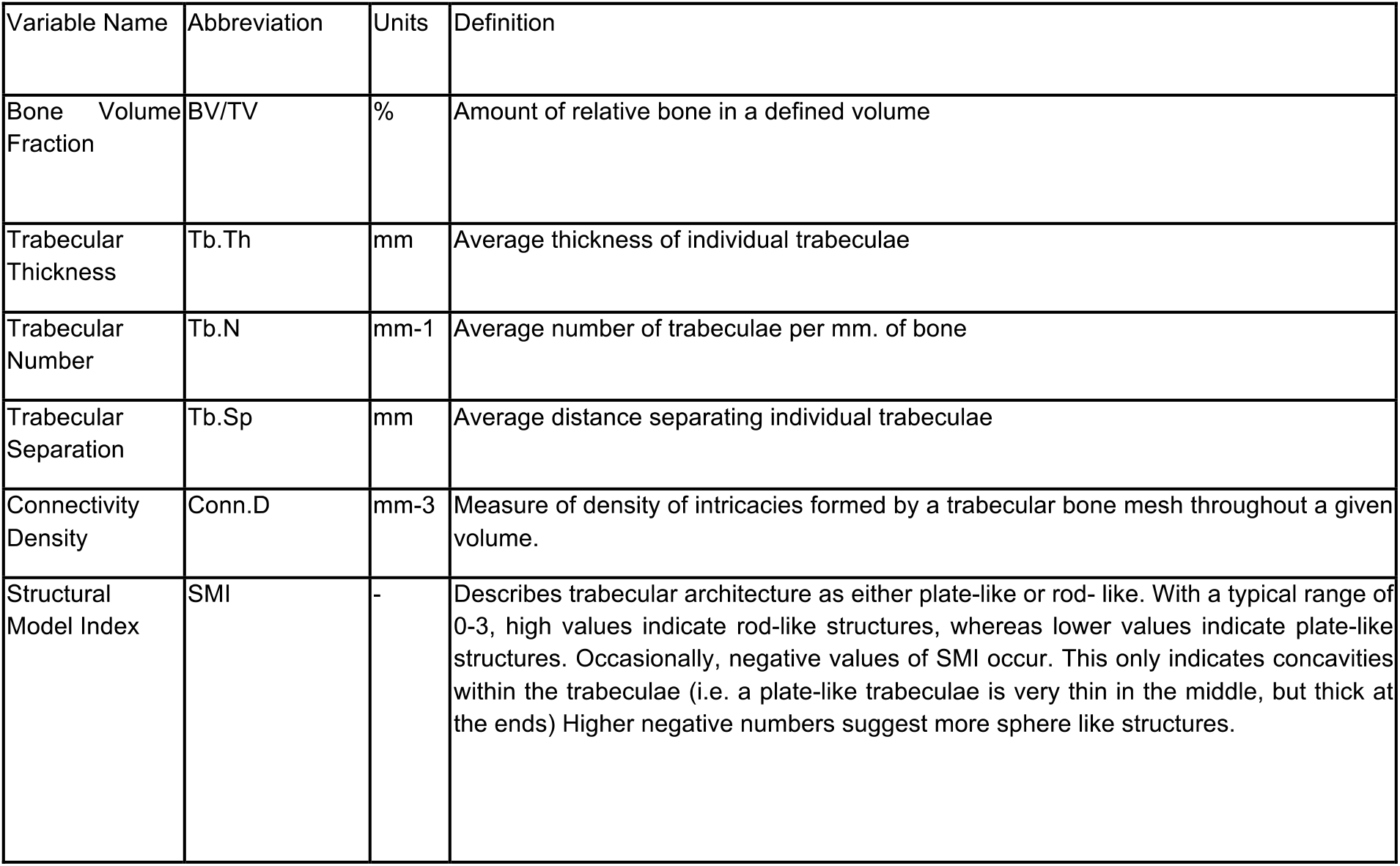

Cortical bone parameters

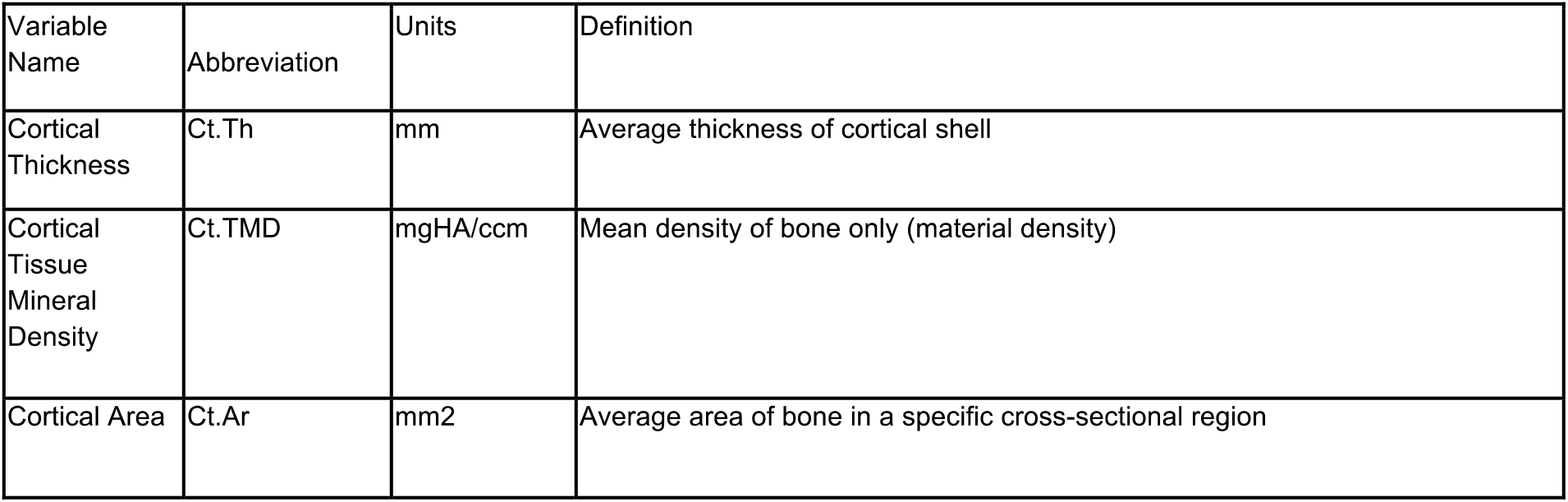
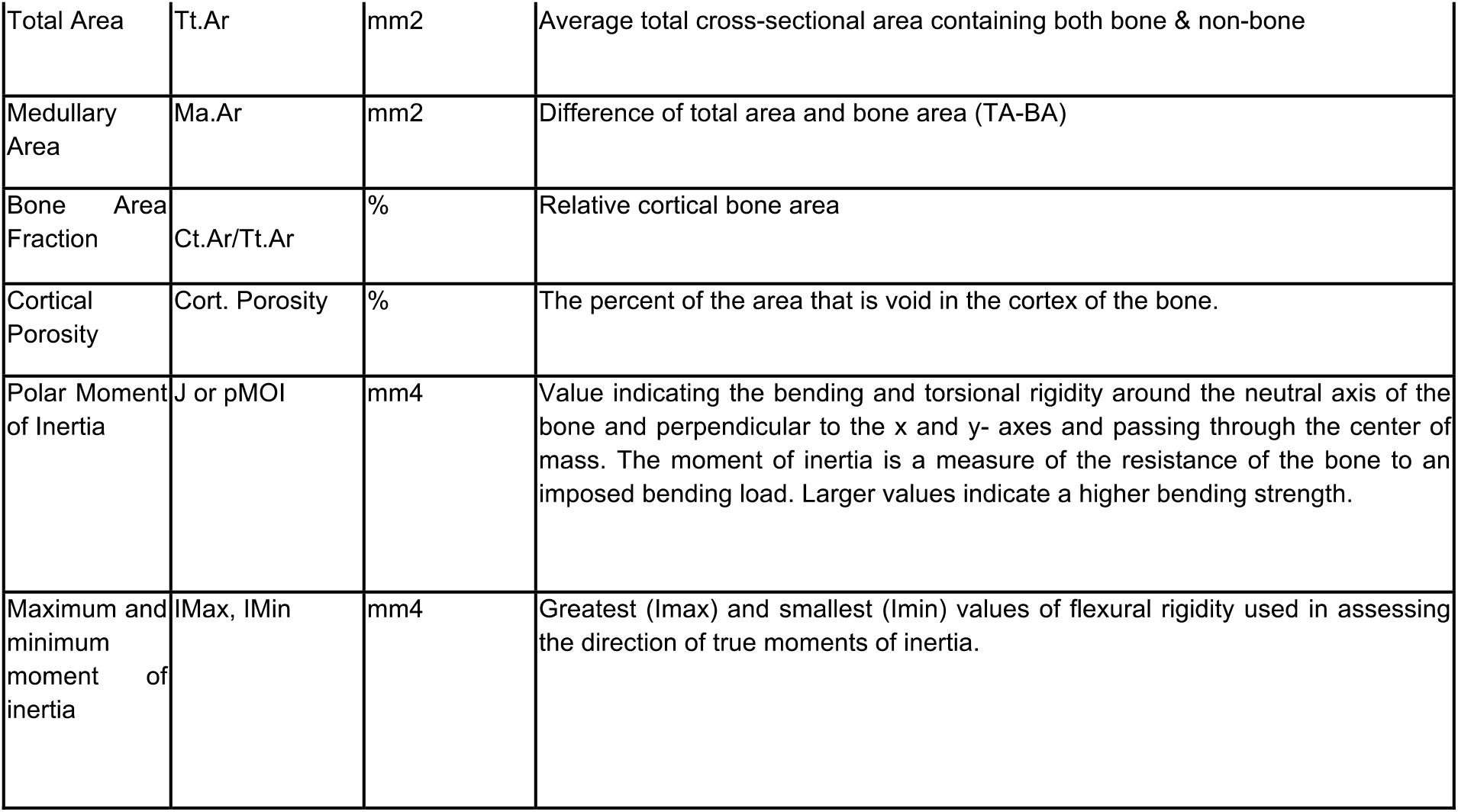

#### Mechanical Testing (Three-point bending)

Femurs were mechanically tested in three-point bending using a materials testing machine (Electroforce 3230, Bose Corporation, Eden Prairie, MN). The bending fixture had a bottom span length of 8 mm. The test was performed with the load point in displacement control moving at a rate of 0.05 mm/sec with force and displacement data collected at 60 Hz. All of the bones were positioned in the same orientation during testing with the cranial surface resting on the supports and being loaded in tension. Bending rigidity (EI, N-mm2), apparent modulus of elasticity (Eapp, MPa), ultimate moment (Mult, N-mm), apparent ultimate stress (oapp, MPa) work to fracture (Wfrac, mJ), and apparent toughness (Uapp, mJ/mm3) were calculated based on the force and displacement data from the tests and the mid-shaft geometry measured with μCT. Work to fracture is the energy that is required to cause the femur to fracture, and it was calculated by finding the area under the force-displacement curve using the Riemann Sum method. Bending rigidity was calculated using the linear portion of the force-displacement curve. The minimum moment of inertia (Imin) was used when calculating the apparent modulus of elasticity.

#### Dynamic histomorphometry

9 days prior to takedown, animals were injected with calcein (20 mg/kg, IP). 2 days prior to takedown, animals were injected with calcein + demeclocycline (10 mg/kg and 40 mg/kg, respectively, IP). Undecalcified vertebra and tibia bone samples were fixed and stored in 70% ethanol. Samples were embedded, then cut by IsoMet 1000 Precision Saw (Buehler, USA) to separate into proximal part of tibia for longitudinal tibia section and distal part of tibia for cortical tibia cross-section. The separation is in the exact midpoint between superior tibiofibular joint and distal tibiofibular joint. The bone samples were dehydrated with acetone and embedded in methyl methacrylate. Consecutive sections were cut in 4 -μm thickness by microtome (RM2255, Leica, Germany). Sections were stained with Von Kossa for showing the mineralized bone and 2% Toluidine Blue (pH3.7) for the analysis of osteoblast, osteoclast and osteoid. The unstained sections were mounted with coverslip for dynamic parameters measurement. The bone sections were viewed with the Nikon E800 microscope equipped with Olympus DP71 digital camera. The image capture was performed by using Olympus CellSens software. The vertebra bone section histomorphometric data was obtained from under 200X magnification in a 1.3 mm x 1.8 mm region away from the growth plate. The tibia longitudinal bone section histomorphometric data was obtained from under 200X magnification in a 0.9 mm x 1.3 mm region away from the growth plate. The tibia cortical bone section histomorphometric data was obtained from under 100X magnification in a 2.6 mm x 1.8 mm region. OsteoMeasure analyzing software (Osteometrics Inc., Decatur, GA, USA) was used to generate and calculate the data. The structural parameters [bone volume (BV/TV), trabecular thickness (Tb.Th), and trabecular number (Tb.N), trabecular separation (Tb.Sp), cortical tissue area (Ct.T.Ar), cortical bone area (Ct.B.Ar), cortical marrow area (Ct.Ma.Ar), cortical bone volume (Ct.BV/TV), cortical thickness (Ct.Th), endocortical perimeter (Ec.Pm) and periosteal perimeter (Ps.Pm) were obtained by taking an average of 2 different measurement from consecutive sections. The parameters were presented according to the standardized nomenclature *(56)*.

### Statistical Analysis

Unless noted otherwise (e.g. RNA-seq analysis), groups were compared using a Mann-Whitney U test (Wilcoxon rank sum test). Where noted, multiple hypothesis testing was accounted for by determining FDR-adjusted q-values (Benjamini-Hochberg method). When sexes were analyzed together (p-combined), data was first analyzed for a significant sex difference in control animals. If a sex difference was not evident, sexes were simply pooled for treatment group analysis. If a sex difference in controls was detected, sex was accounted for in a linear model including an interaction term (value ~ sex*treatment) and the treatment p-value was extracted. P-values for timecourses represent group:day interaction term in a linear mixed model (combined: value ~ treatment*day + sex*treatment + (1|mouse); individual sexes: value~ treatment*day + (1|mouse). All error bars are standard deviation.

## Acknowledgments

The authors thank AbbVie Comparative Medicine for conducting the in-life phase of the 4 month anti-PAPPA mouse study, Lorenzo Benatuil and Tim Esbenshade (AbbVie) for helpful discussions.

## Author contributions

Conceptualization: MM, AF; Methodology: MM, JL, SB, JMT, AB, YK; Investigation: MM, JL, JZS, DH, DB; Project Administration: MLB, RB, YK, AF; Writing: MM, AF; Supervision: AF

## Data and materials availability

All high throughput sequencing data used in this study are available in NCBI GEO (http://www.ncbi.nlm.nih.gov/geo/), accession number GSE144618. anti-PAPP-A antibody and control antibody may be made available subject to a materials transfer agreement with Calico Life Sciences LLC.

---

**Figure S1:**
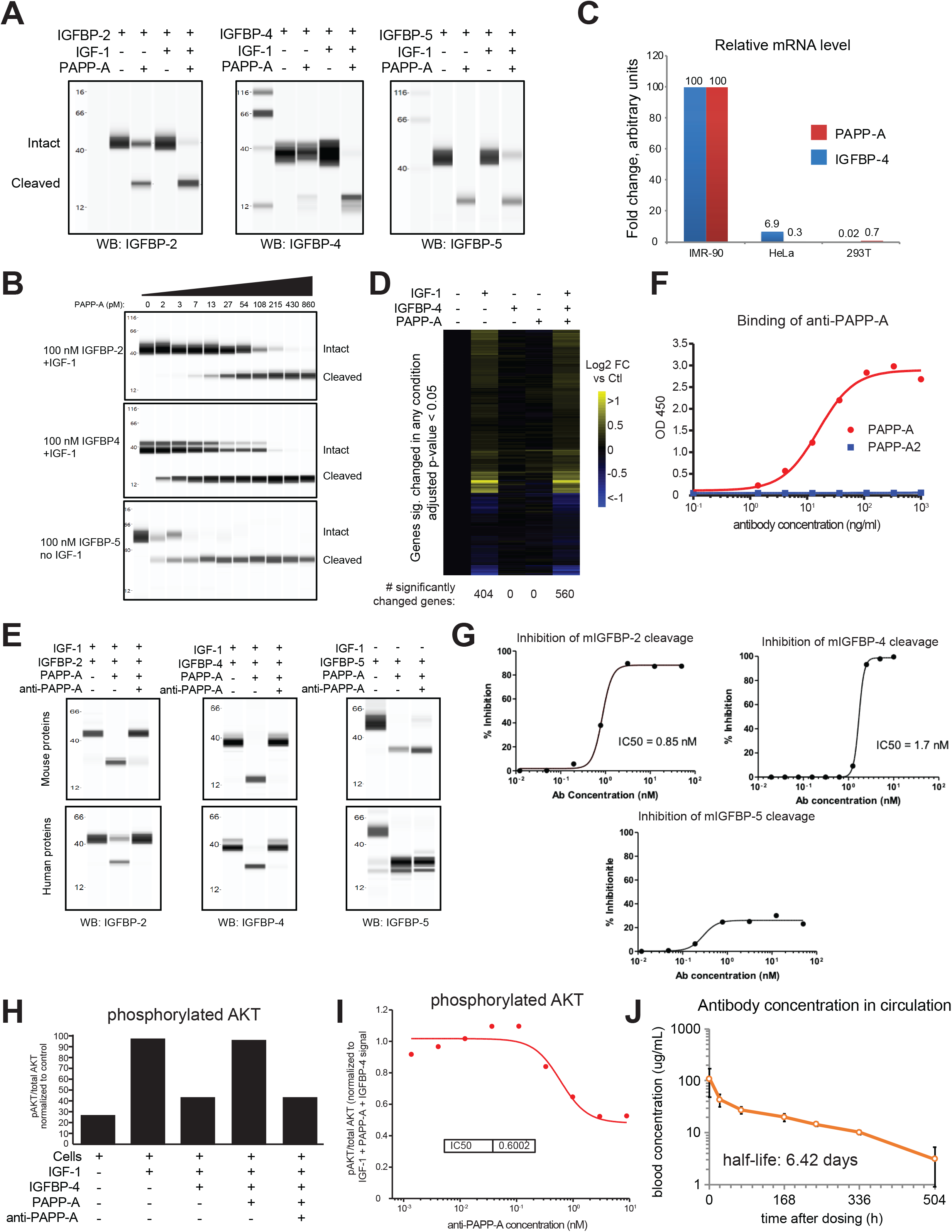
Characterization of a PAPP-A neutralizing antibody. A. Murine PAPP-A cleaves murine IGFBP-2, IGFBP-4, and IGFBP-5. Recombinant proteins were incubated and analyzed for IGFBP cleavage by western blot. B. PAPP-A dose response. Murine PAPP-A cleaves murine IGFBP-2 and IGFBP-4 with similar activity and IGFBP-5 with greater activity. Recombinant proteins were incubated and analyzed for IGFBP cleavage by western blot. C. Cells were serum starved overnight then harvested and processed for RNA. *PAPPA* and *IGFBP4* mRNA were measured by qRT-PCR. n = 3 biological replicates/condition. D. Hela cells were serum starved overnight then recombinant human IGF-1, IGFBP-4, PAPP-A, or the combination thereof was added to the media. Cells were harvested 1 hour later and processed for RNA-seq. n = 3 biological replicates/condition. E. anti-PAPP-A blocks cleavage of human and murine IGFBP-2 and IGFBP-4, but not IGFBP-5. Recombinant proteins were incubated and analyzed for IGFBP cleavage by western blot. F. anti-PAPP-A binds human PAPP-A but not human PAPP-A2. Direct ELISA with the indicated antigens coated on the plate at 1 ug/ml. G. Dose response of anti-PAPP-A inhibition of cleavage of murine IGFBP-2, IGFBP-4, and IGFBP-5. Recombinant proteins were incubated and analyzed for IGFBP cleavage by western blot, then percent inhibition of cleavage was quantified by densitometry. IC50s were calculated for IGFBP-2 and IGFBP-4, but did not reach 50% inhibition for IGFBP-5, so IC50 could not be determined. H. anti-PAPP-A suppresses IGF-1-driven AKT phosphorylation in HEK293 cells. Cells were incubated with recombinant proteins and assayed for pAKT and total AKT 20 min later. I. Titration curve, anti-PAPP-A suppression of IGF-1-driven AKT phosphorylation in HEK293 cells. Signal was normalized to the pAKT/total AKT ratio in the presence of IGF-1, IGFBP-4, and PAPP-A. J. anti-PAPP-A pharmacokinetics indicate an ~6 day half-life. Antibody was IV injected into C57BL/6 mice and blood concentration was monitored by ELISA over the course of 21 days.

**Figure S2:**
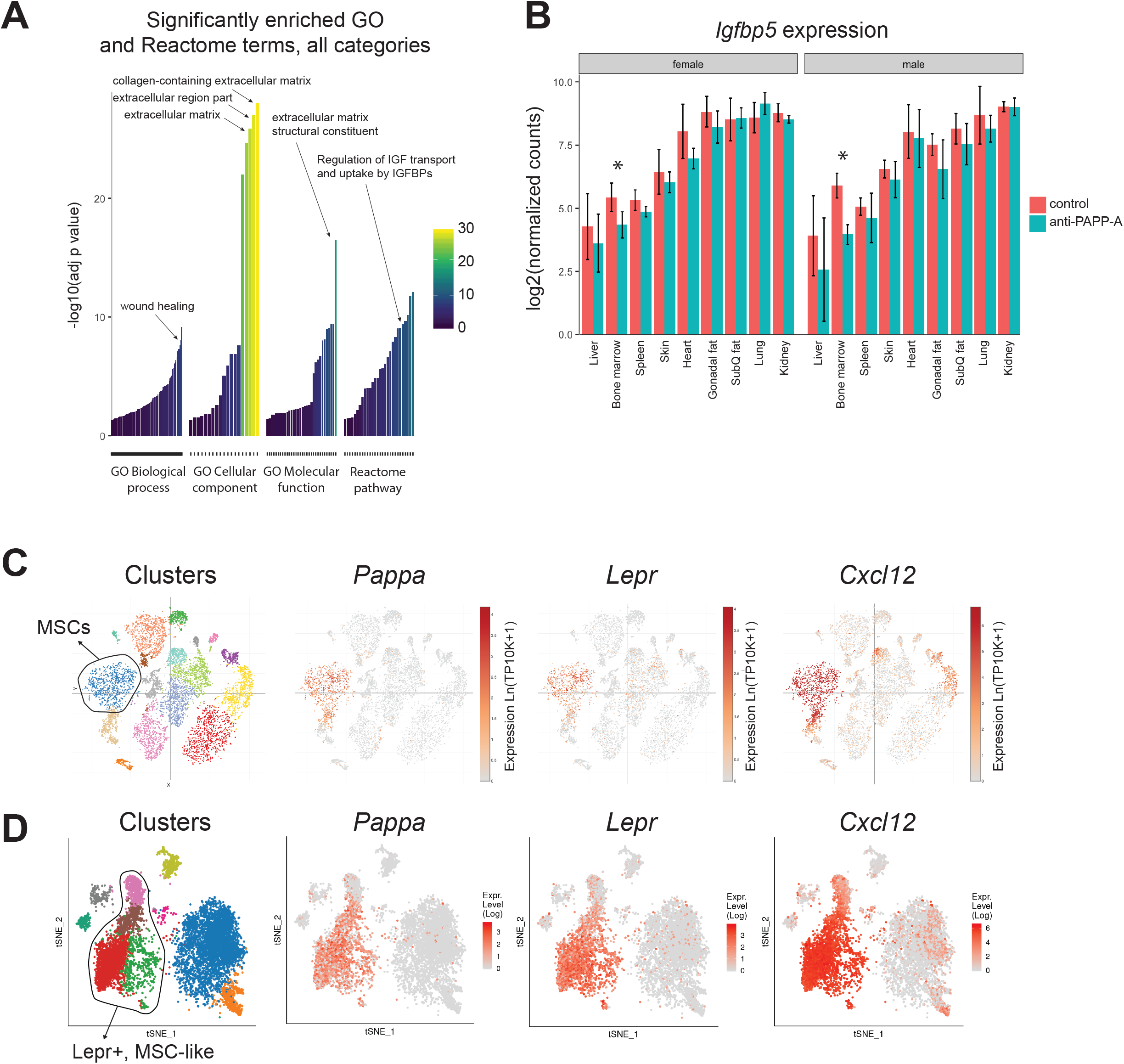
System-wide in vivo evaluation of the effects of anti-PAPP-A identifies reduced collagen-associated extracellular matrix activity and IGF signaling in multiple tissues. A. GO and Reactome terms significantly (adj p < 0.05) enriched in the list of genes differentially expressed by anti-PAPP-A at p < 0.01 (86 genes). B. *lgfbp5* expression level (log2 of normalized counts) in individual tissues, separated by sex. Asterisks identify tissues with a significant (p < 0.05) treatment effect. C. *Pappa* is expressed in MSCs. t-SNE plot depicting scRNA-seq data from mouse bone marrow stroma from Baryawno et al., *Cell*, 2019. *Lepr* and *Cxc/12* are MSC markers. Plots are colored by expression of indicated genes or by post-hoc clustering. Figure adapted from: https://singlecell.broadinstitute.org/singlecell/study/SCP361/. D. *Pappa* is expressed in Lepr+, MSC-like cells. tSNE plot depicting scRNA-seq data from mouse bone marrow stroma from Tikhonova et al., *Nature*, 2019. *Lepr* and *Cxc/12* are MSC markers. Plots are colored by expression of indicated genes or by post-hoc clustering. Figure adapted from: http://aifantislab.com/niche/.

**Figure S3:**
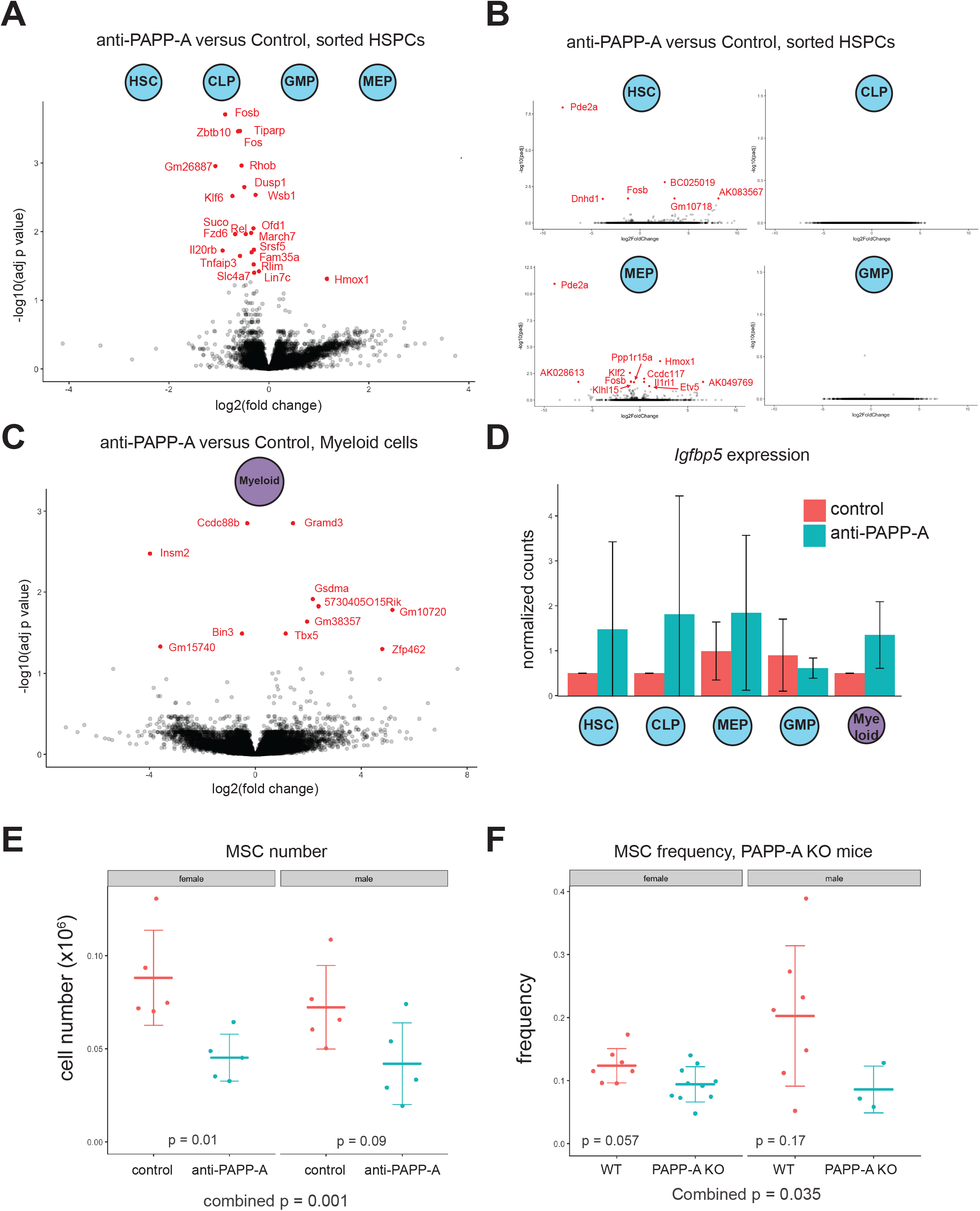
Bone marrow mesenchymal stromal cells respond to anti-PAPP-A. A. Differential expression due to anti-PAPP-A across HSPC populations (HSC, CLP, GMP, MEP). Red dots identify genes differentially expressed at FDR < 0.05. B. Differential expression due to anti-PAPP-A in individual HSPC sub-populations (HSC, CLP, GMP, MEP). Red dots identify genes differentially expressed at FDR< 0.05. C. Differential expression due to anti-PAPP-A in myeloid cells. Red dots identify genes differentially expressed at FDR < 0.05. D. *lgfbp5* expression level (normalized counts) in individual cell types. No significant treatment effects. E. anti-PAPP-A reduces MSC cell number in bone marrow after 4 weeks of treatment (n = 5/sex/group). F. PAPP-A KO mice at 3-6 months of age have reduced MSC frequency in bone marrow (n = 3-10/sex/group).

**Figure S4:**
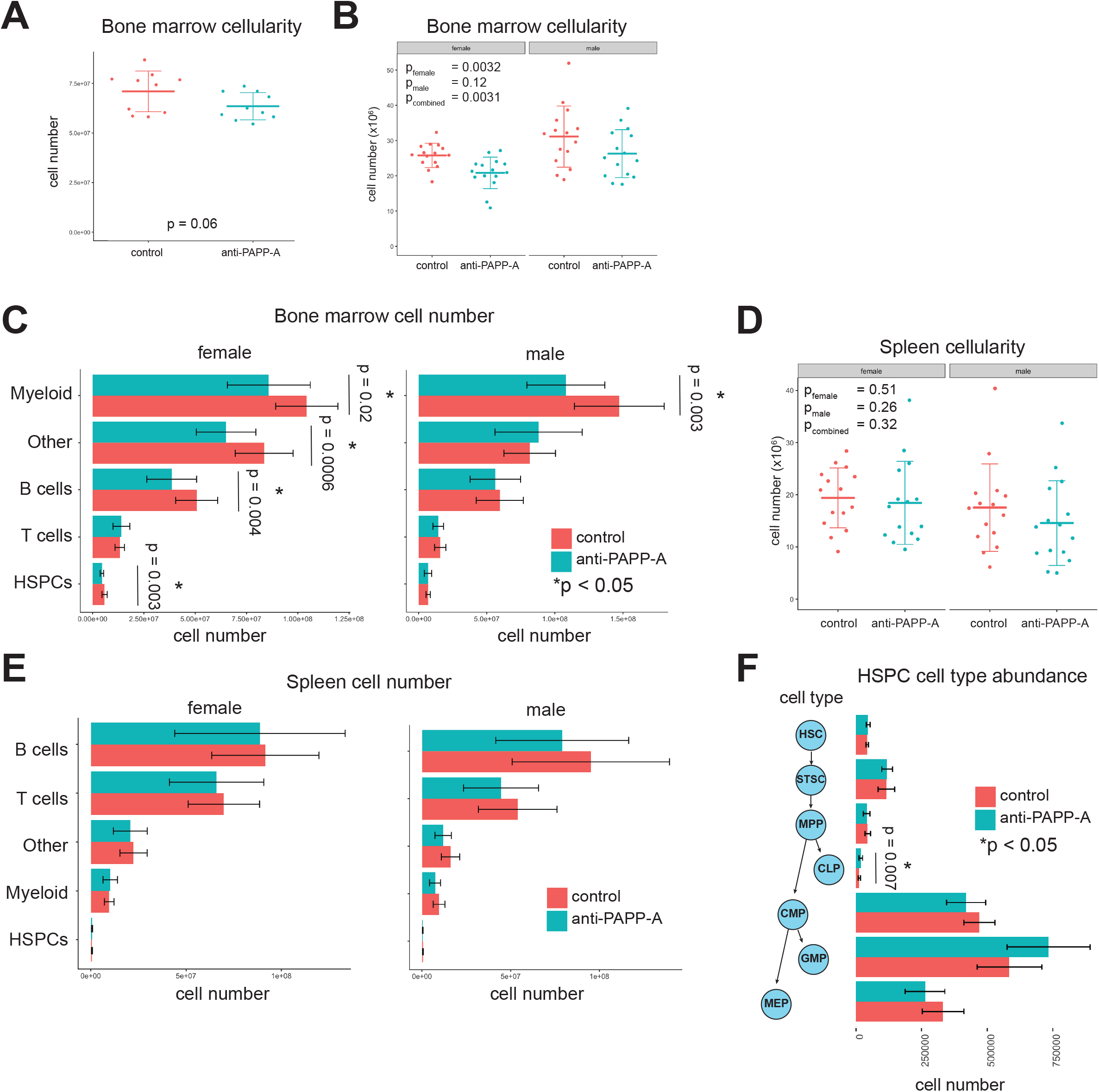
PAPP-A inhibition reduces bone marrow myelopoiesis. A. PAPP-A inhibition for 4 weeks causes a trend toward reduced bone marrow cellularity. Same data as 3B, summed (males only, n = 10/group). B. PAPP-A inhibition for 4 weeks causes a trend toward reduced bone marrow cellularity in both males and females (n = 15/sex/group). C. anti-PAPP-A reduces myeloid cell number inbone marrowin both males and females (n = 15/sex/group). Due to batch effects in this experiment, values have been batch adjusted D. anti-PAPP-A has no effect on spleen cellularity (n = 15/sex/group). E. anti-PAPP-A has no effect on spleen cellularity, analyzed by cell category (n=15/sex/group). Due to batch effects in this experiment, values have been batch adjusted. No significant treatment effects. F. PAPP-A inhibition has minimal effect on HSPCs (males only, n = 10/group).

**Figure S5:**
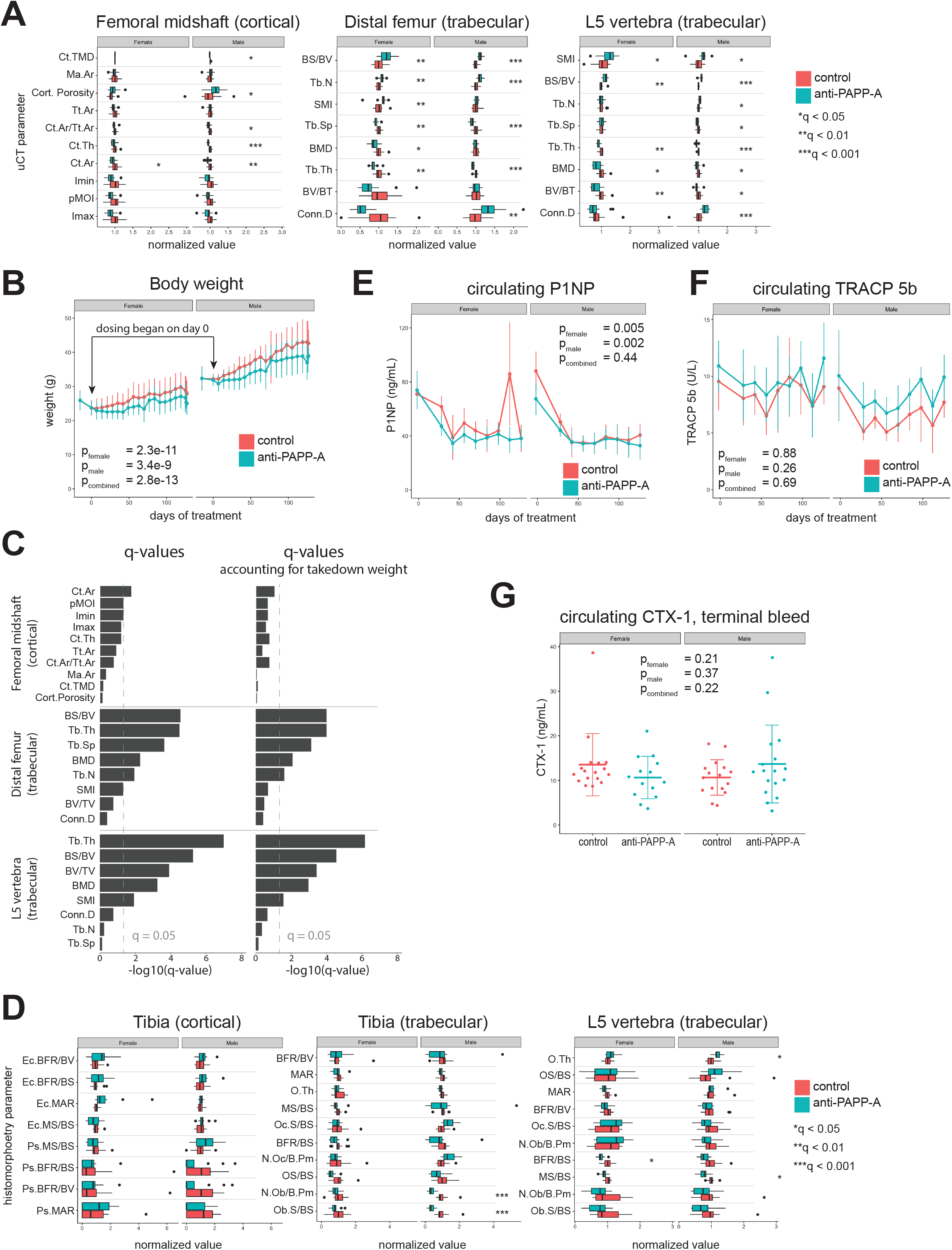
PAPP-A inhibition reduces osteogenesis via reduced osteoblast activity. A. Effect of PAPP-A inhibition on all µCT parameters from femur and vertebra B. PAPP-A inhibition reduces weight gain over time. Dosing began at day 0. C. FDR-adjusted p-values (q-values) for anti-PAPP-A effect on µCT parameters from femur and vertebra, either without (left) or with (right) accounting for takedown weight. Sexes combined. Gray dashed line indicates q = 0.05. D. all dynamic histomorphometry parameters from tibia and vertebra. E. Effect of PAPP-A inhibition on circulating P1NP, a marker of bone formation. F. Effect of PAPP-A inhibition on circulating TRACP 5b, a marker of bone resorption. G. Effect of PAPP-A inhibition on circulating CTX-1, a marker of bone resorption, terminal bleed. (n = 14-18/group).

**Figure S6:**
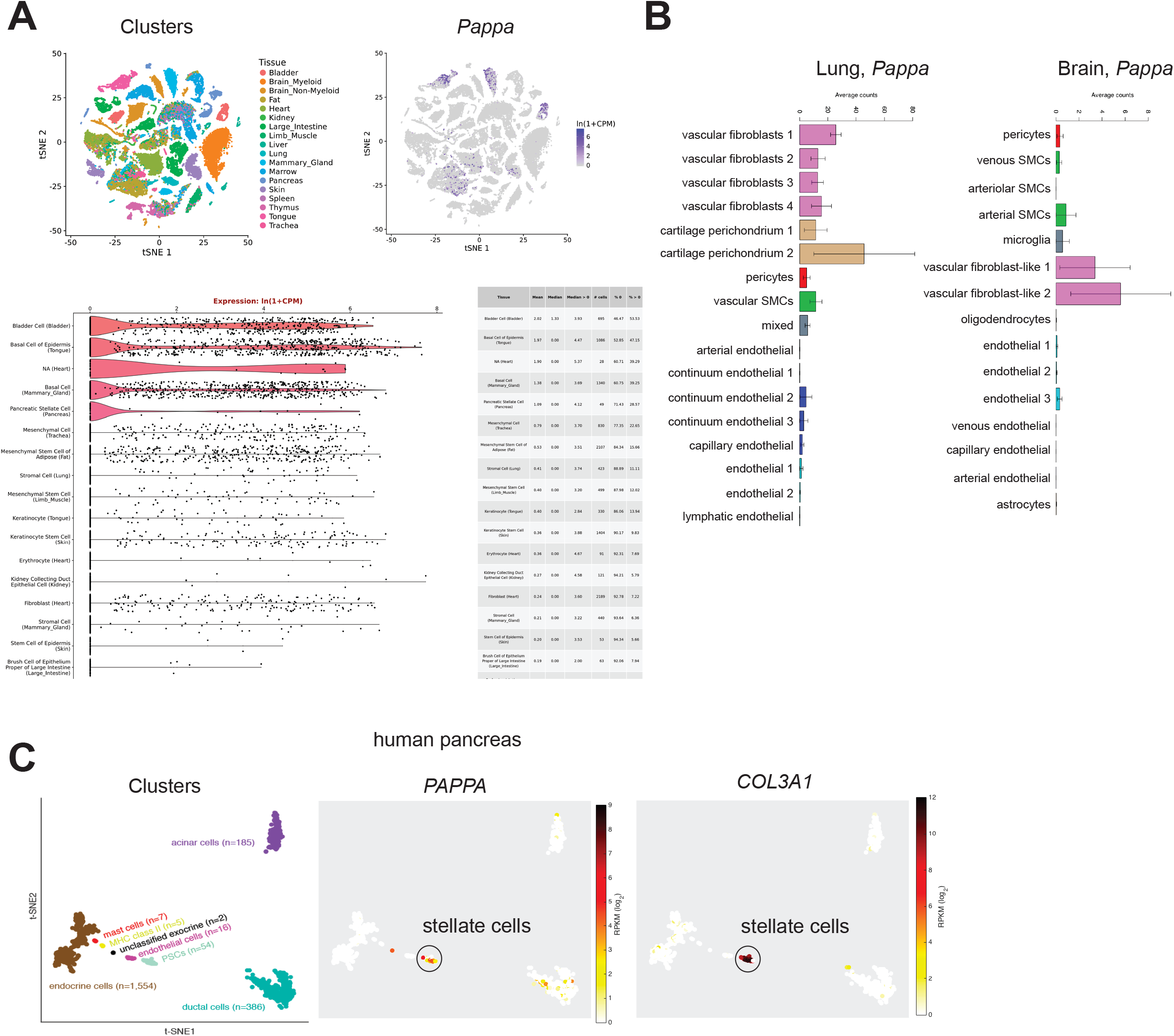
PAPP-A expression in publicly available scRNA-seq datasets. A. scRNA-seq data from mouse tissues, Tabula Muris. *Pappa* is expressed in mesenchymal stem/stromal populations from multiple tissues. Figure adapted from https://tabula-muris.ds.czbiohub.org/. B. scRNA-seq data from mouse brain, Vanlandewijck et al., 2018. *Pappa* is expressed in vascular fibroblast-like cells. Figure adapted from http://betsholtzlab.orgNascularSingleCells/database.html. C. scRNA-seq data from human pancreas, Segerstolpe et al., 2016. *PAPPA* is expressed in stellate cells. Figure adapted from http://sandberg.cmb.ki.se/pancreas/.

**Table S1:** Results of differential expression analysis of all HSPC cell types together, anti-PAPP-A versus Control, 1 week of treatment

**Table S2:** Results of differential expression analysis of individual cell types, anti-PAPP-A versus Control, 1 week of treatment

